# Heat Stress and Soil Microbial Disturbance Influence Soybean Root Metabolite, Microbiome Profiles, and Nodulation

**DOI:** 10.1101/2025.07.13.664636

**Authors:** Dinakaran Elango, Liza Van der Laan, Somayeh Gholizadeh, Maddumage Dona Ginushika Priyadarshani Premarathne, Cole R. Dutter, Cody DePew, Marshall McDaniel, Asheesh K. Singh

**Affiliations:** Department of Agronomy, Iowa State University, Ames, IA 50011, USA; Faculty of Technical Sciences, Department of Agroecology, Aarhus University, Denmark; Department of Ecology, Evolution, and Organismal Biology, Iowa State University, Ames, IA 50011, USA; Department of Plant Science, Penn State University, University Park, PA 16801, USA

**Keywords:** Soybean, Soil microbial disturbance, Heat stress, Root microbiome, Root anatomy, Root metabolite

## Abstract

Heat stress is a major limiting factor for soybean productivity worldwide. Recent studies have highlighted the critical role of the plant microbiome in enhancing plant resilience to heat stress. However, our understanding of the molecular and physiological mechanisms underlying root-microbiome interactions under heat stress remains limited. To elucidate the role of native soil microbes in the heat tolerance of soybean genotypes, we analyzed rhizosphere bacterial and fungal communities via 16S rRNA and ITS sequencing, and characterized root metabolites and anatomical traits in response to microbiome composition and heat stress. Soybean plants were grown under controlled conditions in either natural soil containing native microbiota or in microbiome-disturbed soil (via 3-hour autoclaving), under both optimal and elevated temperature regimes. Alpha and beta diversity analyses revealed significant microbial shifts between treatments. Distinct clustering of bacterial, fungal, and metabolite profiles was observed under high temperature and microbial disturbance. Nodule-forming bacteria such as *Rhizobium* and *Janthinobacterium* were markedly suppressed, and belowground traits exhibited sensitivity, with significantly reduced nodule numbers and nodulation efficiency under high temperature and soil microbial perturbation. Non-targeted root metabolomics identified 372 differentially accumulated metabolites. Integrative multi-omics analysis revealed associations between metagenomic profiles, metabolite levels, and nitrogen-fixation traits, implying a coordinated modulation of root physiological processes. These findings contribute to a growing understanding of how heat stress interacts with rhizosphere microbial communities and may support future efforts in breeding climate-resilient soybean cultivars.

## Introduction

Soil is the largest reservoir of microbial diversity on Earth, holds more than half of the world’s biodiversity [1]. Remarkably, a gram of dry soil may contain between four million to two billion bacterial cells [2]. This immense microbial diversity plays a crucial role in maintaining the health of plants, humans, and animals, either directly or indirectly [3]. The microbial communities associated with these organisms act as an’extended genotype,’ commonly referred to as the’second genome’ of the organism [4–6]. These interconnected microbiomes significantly influence one another’s well-being [7, 8], creating a delicate balance that is essential for overall health. However, this balance is increasingly threatened by climate change, which directly impacts the health of plants, soils, and microbiomes [9–11]. Changes such as increased temperature stress, water-limiting conditions, altered soil pH levels, and shifts in organic matter content lead to fluctuations in microbial prevalence [12]. Consequently, understanding the interactions between soil microbes and plants under extreme climatic conditions is critical for developing climate-resilient crops and improving soil health, both of which are essential for ensuring sustainable food security [13].

Microbial communities in healthy soils play a crucial role in maintaining plant health by enhancing nutrient cycling, pathogen suppression, and stress resilience [14]. For instance, *Rhizobium* spp. engage in a symbiotic relationship with soybean roots, converting atmospheric nitrogen into usable forms for the plant [15]. Emerging evidence suggests that this relationship might also confer heat stress tolerance to soybeans [16]. Additionally, some microbes aid plants in accumulating osmoprotectants, such as proline and trehalose, which help maintain cell turgor and protect cellular structures under heat stress [17]. Specific endophytic bacteria in soybean have been found to enhance heat tolerance by modulating the plant’s antioxidant defense system, thereby reducing oxidative damage during heat stress [18]. Therefore, microbes that are beneficial in alleviating plant stresses hold promise for improving plants adaptability to changing climatic conditions.

Soil microbes also play a critical role in modulating plants responses to invading pathogens. Extensins, cell-wall glycoproteins in plant roots, undergo extensive modifications when pathogenic microbes are detected, strengthening the plant cell wall and thereby restricting pathogen entry [19, 20]. Additionally, extensin molecules, together with root-tip border cells, form the’root extracellular trap,’ a defense mechanism secreted into the rhizospheric soils to combat invading pathogens. This mechanism works by interacting with pectins and arabinogalactan proteins, weakening the attachment of pathogenic microbes to the roots [21–23].

Furthermore, soil microbes are involved in root-to-shoot signaling via root exudates, which modulate induced systemic resistance (ISR) in plants. This highlights how soil microbes contribute to crop resilience [24]. For instance, microbes isolated from the roots of foxtail millet, such as *Pseudomonas fluorescens, Enterobacter hormaechei,* and *Pseudomonas migulae,* have demonstrated enhanced performance under drought stress conditions [25]. With advances in metagenomics and synthetic biology, there is potential to engineer or condition the soybean microbiome to further enhance its resilience to heat or drought stress [26].

The interactions between soybean and its associated microbes provide a promising avenue for enhancing the crops tolerance to heat stress. As global temperatures continue to rise, leveraging these interactions through targeted microbiome engineering or conditioning could be a pivotal strategy for improving soybean’s heat stress tolerance. Soybean (*Glycine max*), a globally significant legume crop, is particularly susceptible to heat stress. For every degree Celsius increase above the optimum temperature, there is approximately a 17% yield decline in soybean [27]. Soybean is one of the most important crops in Iowa [28, 29], where farmers cultivate soybeans primarily in two major soil series: Clarion and Sparta [30]. The Clarion soil series has a clay loam texture with moderate to high organic matter, while the Sparta series, with its loamy sand texture, has low organic matter and is prone to soybean cyst nematode (SCN) issues. In our experiment, we used the Clarion soil series to investigate the diversity of soil microbes and their role in mitigating heat stress in soybean.

Our main goal of the experiment is to understand microbial abundance and its prevalence with respect to high-temperature stress through amplicon sequencing (16S and ITS) and to explore the role of these microbes in stress tolerance through non-targeted global metabolomic profiling and root anatomical studies. Understanding the intricate relationships between soil microbes and plants under high-temperature stress will pave the way forward, providing crucial feedback loops to breeders in developing heat-tolerant cultivars suited for future climatic conditions.

## Materials and Methods

### Soil sample collection and characterization

We used clarion soil series with loamy texture (45% sand, 33% silt, and 22% clay) from the Iowa State University’s (ISU) agronomy research farm in Ames, USA (42°01’05.5“N 93°46’11.2”W). Topsoil (0-20 cm) was taken after the harvest of soybean. After removal of roots and debris, the soil was sieved through a 2 mm mesh, homogenized by mixing and divided into two portions. One portion of the soil was stored in a cold room (4-5 °C) for approximately 15 days and labeled as non-autoclaved soil with the natural microbiome. The other portion was kept in the same conditions, but autoclaved at 121 °C and 15 psi for 3 hrs one day prior to planting resulting in the autoclaved soil with disturbed microbiome. This procedure was done to reduce soil microbial biomass. Both autoclaved and non-autoclaved soils were mixed with sterile sand (1:1 v/v) to get better drainage and also to sample the roots at the end of the experiment. The physical properties of the autoclaved and non-autoclaved soil samples were determined using a laser diffraction spectroscopy at ISU, Ames [31]. Soil physical properties did not differ significantly between autoclaved and non-autoclaved treatments, as shown in Table S1. Soil samples were further sent to Mid-west labs for chemical characterization. Soil chemical properties were analyzed and are presented in Table S2 and S3. Soil microbial respiration (CO_2_) for both autoclaved and non-autoclaved soil samples were determined using an infrared gas analyzer (Licor 500, Lincoln, NE USA). A three-day incubation experiment revealed a significant difference in net ecosystem exchange (NEE) of CO₂ between the two treatments (p < 0.05), with autoclaved soils showing reduced NEE compared to non-autoclaved soils (Fig. S1).

### Plant materials and experimental conditions

Four soybean genotypes were used including PI 639693, a heat-tolerant cultivar, PI 89008, a heat susceptible cultivar, IAS19C3, an elite cultivar from the ISU soybean breeding program, and the standard reference genotype Williams 82. In this study, we investigated how soil microbiome disturbance influences the response of soybean genotypes to heat stress. Therefore, plants grown in non-autoclaved and autoclaved soils holding its natural or disturbed microbial community were subjected to different levels of heat stress. All the seeds were then surface sterilized using 6.25% sodium hypochlorite solution for 45 seconds, followed by three rinses with deionized distilled water [32]. Surface-sterilized seeds were planted in sterile pots (1 gallon) filled with autoclaved and non-autoclaved soils. All pots were watered daily with a shower of distilled water as an accessible proxy for rainwater that avoids chlorine and other tap water additives. After seed germination, only one healthy seedling was kept until sampling time. Pots were spatially randomized according to the treatments in four different Enviratron growth chambers, providing long days of 16 h light and 8 h dark, light intensity 500 µE m^−2^ s^−1^, relative humidity 60%, and CO_2_ 400 ppm [33]. Heat treatments were maintained for the growth chambers at either a high temperature at 38/28 °C (day/night) or at 28/21 °C (day/night) for an optimum temperature. We thereby had four treatments including high temperature with non-autoclaved soil (NH), high temperature with autoclaved soil (AH), optimum temperature with non-autoclaved soil (NO), and optimum temperature with autoclaved soil (AO). All samples were harvested at the V_5_ stage for every four replications of the genotypes to evaluate different analyses.

### Rhizosphere microbial community profiling

After removing the plant from each pot, the excess soil was manually shaken from the roots, leaving approximately the fine soil particles still attached to the roots. We then placed the roots in a 50 ml sterile Falcon tube with 30 ml of sterile ultra-pure water to separate the remaining soil (rhizosphere) from the roots. The soil cleaned from the roots was then spun at 15000 g for 15 min to form tight pellets, from which all supernatant was removed, snap-frozen, and stored at-80 °C until DNA extraction [34]. DNA was extracted from rhizosphere soil samples using Qiagen’s DNeasy power soil kit, according to the manufacturer’s instruction, and stored at-20 °C prior to Illumina MiSeq library preparation. For bacterial community characterization, the V_4_ hypervariable region of the bacterial 16S rRNA gene was amplified with the primers 515F (5ʹ-GTGYCAGCMGCCGCGGTAA-3ʹ) and 806R (5ʹ-GGACTACNVGGGTWTCTAAT-3ʹ) [35, 36]. For fungal community characterization, the ITS1–ITS2 region of the fungal ITS gene was amplified with the primers ITS1F (5ʹ-CTTGGTCATTTAGAGGAAGTAA-3ʹ) and ITS2 (5ʹ-GCTGCGTTCTTCATCGATGC-3ʹ) [37]. Library preparation and amplicon sequencing was performed on Illumina Miseq (Illumina, USA) at ISU’s DNA facility.

### Microbial data processing

Bacterial and fungal sequences were analyzed using the DADA2 pipeline (v. 1.30.0) [38]. For fungi, the primers were first trimmed off the demultiplexed sequences using Cutadapt version 4.8 [39]. Low-quality reads were filtered using filterAndTrim function with the following parameters (maxN = 0, maxEE = c(5, 5), truncQ = 2, minLen = 50, rm.phix = TRUE, compress = TRUE) and then error rates were estimated using the learnErrors function. The denoised sequence table (ITS sequence variants) was constructed after sample inference and merging paired reads. The bacterial raw demultiplexed reads were also processed using the DADA2 pipeline to denoise (with the following parameters used in the filterAndTrim-step: maxN = 0, maxEE = c(2,2), truncQ = 2, truncLen=c(250, 220), trimLeft=20, rm.phix = TRUE, compress = TRUE), dereplicate reads, merge pair-end reads and remove chimeras. Taxonomic affiliation was then performed with DECIPHER version 2.30.0 [40] based on the IDTAXA algorithm and the SILVA version 138.1 [41] as a reference database for fungal and bacterial sequences, respectively. To construct the phylogenetic tree, we first performed a sequence alignment (AlignSeqs function) of all the fungal and bacterial amplicon sequence variants (ASVs) using the DECIPHER R package version 2.30.0 [40] and constructed a neighbor-joining (NJ) tree in the PHANGORN R package version 2.11.1 [42]; next, we fitted a maximum likelihood tree (GTR model) using the NJ tree as a starting point in DADA2. Finally, the ASV abundance, taxonomy table, and phylogenetic tree were combined with the sample metadata to create a phyloseq object with 3176 and 2468 taxa assigned to seven taxonomic ranks for bacteria and fungi, respectively. All singletons and all reads from other than bacterial or fungal origin (i.e. mitochondria, chloroplasts, and protists) were removed from the datasets. To account for large differences in read numbers, all samples with less than 11000 and 2000 reads (respectively in 16S and ITS data) were removed which resulted in the removal of one sample for bacteria and 33 samples for fungal dataset. Furthermore, all ASVs with relative abundance of <0.005% were removed from both datasets. Log transformation was used to normalize the datasets.

### Statistical analysis of microbiome

Alpha diversity was calculated via the Shannon diversity (H‘) and observed operational taxonomic unit (OTU) richness indices using the package Phyloseq in R [43]. Parametric assumptions were verified using the Shapiro-Wilk normality test in the R base. Statistical analysis for alpha diversity was also done with the function “Kruskal. test” or “pairwise.Wilcox.test” in the R base to compare alpha-diversity values among non-parametric samples. ANOVA variance with Tukey comparison tests were used for parametric samples. For beta-diversity analyses, OTU tables were normalized by the log transformation method using the R package DESeq2 [44]. Bray-Curtis distance, as well as UniFrac distances (weighted and unweighted), were calculated from the normalized OTU tables using the “ordinate” function of the R package Vegan [45]. We conducted principal coordinates analysis (PCoA) with UniFrac matrics (weighted and unweighted) at the ASV level using the Phyloseq R package. Permutational multivariate analysis of variance (PERMANOVA) was coupled with dispersion analyses performed with the adonis and betadisper functions, respectively, using the ‘vegan’ R package. The information of beta and alpha diversity considering all factors and their interactions were listed in Table S4 and S5. Venn diagrams were constructed to visualize the proportion of groups exclusive and shared between treatments using R package microeco [46]. To identify individual microbia differentially recruited across different treatments and soybean genotypes, we implemented a differential analysis of the count data using negative binomial generalized linear models using the DESeq2 package in R with default settings. The linear discriminant analysis effect size (LEfSe) was also used to identify the significant biomarkers in microbial taxa among different treatments [47] and their visualization using microeco packag in R [48]. The R package microbiomeutilities [49] was used for exploring the soybean rhizosphere core microbiome and the abundance-prevalence relationship analyses identified at a 50% prevalence threshold. Redundancy analysis (RDA) was used to evaluate the relationships between plant phenotypic attributes and rhizosphere ASVs. Mantel test was used to study the relationship between alpha diversity and plant metabolites and between the enriched ASVs and environmental factors. Mantel tests, principal coordinate analysis (PCoA), and RDA were performed using the “vegan” package in R v. 3.6.3 [50]. Data visualization was performed with the R package ggplot2 [51]. Tax4Fun2 (v1.0) was further implemented in default settings to predict and compare the treatment-specific functional pathways and functional redundancy of bacterial communities [52]. Ecological guilds of fungal ASVs were also predicted using FUNGuild [53].

### Non-targeted root metabolite profiling

Root metabolome was extracted using ultra-pure water and the extract was lyophilized using the freeze-drier. The lyophilized samples were then sent to Metabolon for global non-targeted metabolite profiling. Samples were processed using the automated MicroLab STAR system (Hamilton Company), with recovery standards added for quality control. Proteins were precipitated using methanol under vigorous shaking (2 min, GenoGrinder), followed by centrifugation. The resulting extract was divided into five fractions: four for UPLC-MS/MS analysis under distinct chromatographic and ionization conditions, and one reserved as backup. After solvent removal (TurboVap), dried extracts were stored under nitrogen overnight. Prior to analysis, samples were reconstituted in method-specific solvents containing internal standards to ensure consistency. Analyses were performed using a Waters ACQUITY UPLC system coupled to a Thermo Q-Exactive mass spectrometer with a HESI-II source and Orbitrap analyzer (35,000 resolution). The methods included reverse-phase (RP) UPLC-MS/MS in positive ion mode optimized separately for hydrophilic and hydrophobic compounds using a BEH C18 column, RP in negative ion mode with ammonium bicarbonate (pH 8), and HILIC in negative ion mode using a BEH Amide column with ammonium formate (pH 10.8). Mass spectra were acquired in MS and data-dependent MSⁿ modes with dynamic exclusion, covering a scan range of m/z 70–1000. Quality control measures included pooled technical replicates, blanks, and spiked standards to assess instrument performance and process variability via relative standard deviation (RSD).

### Data extraction and compound identification

Raw data were processed, peak-identified, and quality controlled using Metabolon’s proprietary software and hardware platforms. Compound identification was achieved by matching retention index (RI), accurate mass (±10 ppm), and MS/MS spectral data to a library of over 3,300 authenticated standards and recurrent unknowns. MS/MS scores were calculated based on spectral similarity to reference compounds. Data curation removed artifacts and ensured consistent identification. Peak areas were quantified using area-under-the-curve. For multi-day runs, data were normalized within each run-day block to correct for inter-day instrument variability. Additional normalization to protein content or other sample-specific factors was applied as needed.

### Metabolite data processing and bioinformatics

To the identification of potentially significant marker metabolites, the differential metabolites were screened based on the combination of fold change and variable importance in project (VIP) value of the partial least squares discriminant analysis (PLS-DA) model (fold change ≥ 1.5, VIP ≥ 2). PLS-DA was carried out on log-transformed and auto-scaled (i.e. mean-centered and divided by the standard deviation of each feature) metabolite concentrations using MetaboAnalyst [54]. Pearson correlations between plant microbiome and metabolites were examined in R and heatmaps were drawn using R package microeco [46]. Finally, Euclidean distance measure and Bray-Curtis clustering algorithm was used to measure the difference in the VIP metabolites between the treatments. For that, a linear regression model was fit to the data, the R^2^ statistics calculated, and then this correlation was tested for significance using both r and n, all with type = “lm” in R package microeco [46].

### Root anatomical traits profiling

The taproot samples (5 cm long) were collected and immersed upon excising from the plant using 75% ethanol. Samples were then used for dissection to study the anatomical features of different soybean genotypes using the Laser Ablation Tomography (LAT) technique at Penn State, University Park, PA, USA [55]. For root anatomy, several taproots of each genotype were gently dissected and divided with a razor blade into 5 cm lengths and dried using an automated critical point dryer (CPD) according to the manufacturer protocol, as previously described. Segments of dried taproots were ablated by a laser beam (Avia 7000, 355 nm pulsed laser) to vaporize the root at the camera focal plane ahead of an imaging stage, then cross-section images were taken using a Canon T3i camera with a 5x microlens (MP-E 65 mm) on the laser-illuminated surface. The resulting images were analyzed by the software MIPAR [56] to obtain areas of the cross-section, stele, and metaxylem vessels. The theoretical axial water conductance of taproots was calculated according to Tyree and Ewers [57]. Picasa software was used to convert the semi-monochromatic high-quality LAT images into complementary dyadic colored images with a bright white background using the following steps: a) Heat map the laser image to 0% hue and nearly 50% fade, b) invert the colors, c) cross process, then finally d) Orton-ish by 0% bloom, almost 50% brightness and 0% fade, leading to qualitatively distinguish the secondary cell wall elements. The pith and xylem tissues were extensively mapped with respect to different treatments.

### Measurement of nodules and leghemoglobin content

The number of nodules present in the tap and secondary roots was counted and weighed manually. Then the root nodules were flash-frozen and stored at-80 °C until further analysis. The leghemoglobin was extracted using Drabkin’s reagent, and pure hemoglobin was used to make a standard curve [58]. To extract leghemoglobin, 0.5 g of nodule tissue was ground using a small mortar and pestle until fully homogenized. The ground tissue was then transferred to a 50 mL centrifuge tube, and 3 mL of Drabkin’s reagent was added. The tube was sealed tightly and placed on a shaker, where the mixture was gently agitated for 15 minutes. The homogenized solution was centrifuged at 500 x g for 15 minutes. After centrifugation, the supernatant was carefully transferred to a 10 mL volumetric flask, ensuring no solid material was transferred. Another 3 mL of Drabkin’s reagent was added to the centrifuge tube containing the residual nodule tissue, which was agitated using a small stirring rod until the particles were evenly dispersed. The mixture was centrifuged again at 500 x g for 15 minutes, and the supernatant was transferred to the same volumetric flask. This process was repeated once more to ensure maximal extraction of leghemoglobin. The combined supernatants were brought to a final volume of 10 mL with Drabkin’s reagent. The solution was then transferred to a fresh centrifuge tube and centrifuged at 20,000 x g for 30 minutes to remove any remaining particulates. The final leghemoglobin solution was stable for several hours and was used for subsequent analysis.

To prepare the hemoglobin stock solution, 1800 mg of powdered hemoglobin was transferred into a 10 mL volumetric flask using a funnel. Any remaining powder on the funnel and weigh boat was washed into the flask with Drabkin’s reagent to ensure full recovery of hemoglobin. The flask was then made to volume with Drabkin’s reagent and mixed thoroughly by inverting 8-10 times until the powder was completely dissolved. The stock solution was stored in a light-impermeable bottle at 2-8°C and used within 30 days. A working standard solution was prepared by pipetting 40 µL of the stock solution into a 10 mL volumetric flask and making up to volume with Drabkin’s reagent. This solution was mixed thoroughly by inversion. Subsequently, working standards were prepared for 0, 60, 120, and 180 mg ml^-1^. The resulting solutions were mixed thoroughly by inverting each tube 8-10 times to ensure homogeneity. The absorbance of the standards was measured at 540 nm using a spectrophotometer. A calibration curve was generated, which was linear and passed through the origin.

### Data analysis of root phenes

Effects of experimental treatments on all root anatomical traits including shoot fresh weight (g), number of nodules per plant, nodule weight per plant (g), nodulation efficiency, total area (mm^2^), pith area (mm^2^), xylem area (mm^2^), and percent pith (%) were evaluated by analysis of variance (ANOVA) using the Student’s t-test in R base (Version 4.3.2). RDA were conducted using the Vegan package in R [45]. Pearson pair-wise correlations were used to examine the relations between plant phenotypic parameters and plant microbiome. The differential significance analysis of nodule and anatomical variables and its visualization were performed using R package microeco [46].

## Results

### Rhizosphere microbial diversity, community structure, and composition responses to heat stress and soil biotic disturbance

After quality and filtering, we sequenced 2,411,676 bacterial reads across 64 samples (one sample was removed from statistical analyses due to a low read depth < 2500) with a minimum read depth of 13,806 and a maximum of 52,213. After dereplicating, denoising, and removing chimeras, the number of 2,193,262 high-quality sequences was obtained that clustered into 3176 ASVs at 99% similarity. For ITS, we sequenced 2,659,583 raw reads across 64 samples (33 sample were removed from statistical analyses due to low read depths < 2000) with a minimum read depth of 2263 and a maximum of 134,050. The amount of 1,863,065 high-quality sequences clustered into 2468 unique taxa of fungi with a similarity cutoff ≤ 97% was acquired after dereplicationg, denoising, and removing chimeras. After discarding uncharacterized, low-abundance, and spurious data, 2569 and 2468 ASVs were identified for bacteria and fungi, respectively. All ASVs with relative abundance of <0.005% were then removed from the datasets. In the end, we detected total of 1441 ASVs for bacteria and 1023 phylotypes of fungi across samples.

### Bacteria

PCoA analyses generated using phylogenetic relationships with both weighted UniFrac distance matrix, sensitive to relative abundances of taxa, and unweighted UniFrac distance, sensitive to unique taxa, confirmed an observed discrimination of rhizobacterial communities across different treatment combinations (Fig. 1A and B). PCoA ordination plots declared that the soil type (autoclaved vs. non_autoclaved) explained 51.7% and 36% of the variance of treatment and the temperature (high vs. optimum) could explain 23.2% and 19.8% of the overall variance of the bacterial data set on weighted and unweighted Unifrac distances, respectively. PCoA results also confirmed that the samples were derived from non-autoclaved soil generally clustered close to each other at both high and optimum temperatures, while autoclaved soil with microbiome disturbance exhibited a clear clustering in both thermal stress and far from those of non-autoclaved soils as depicted in Fig. 1A and B. PERMANOVA based on the distance matrices (Adonis) also displayed a significant contribution of different treatments (R^2^ = 76.75%, P = 0.001, weighted; R^2^ = 63.55%, P = 0.001, unweighted; Table S2), and genotype-by-treatment interactions (R^2^ = 82.16%, P = 0.001, weighted; R^2^ = 71.7%, P = 0.001, unweighted; Table S4). Further, PERMANOVA showed that rhizobacterial communities of the genotypes used in this study accounted for very small, but not significant of the overall variation between bacterial communities and indicated a consistent diversity pattern across different genotypes (R^2^ = 1.55%, P = 0.984, weighted; R2 = 2.31%, P = 0.987, unweighted; Table S4).

**Figure 1.**
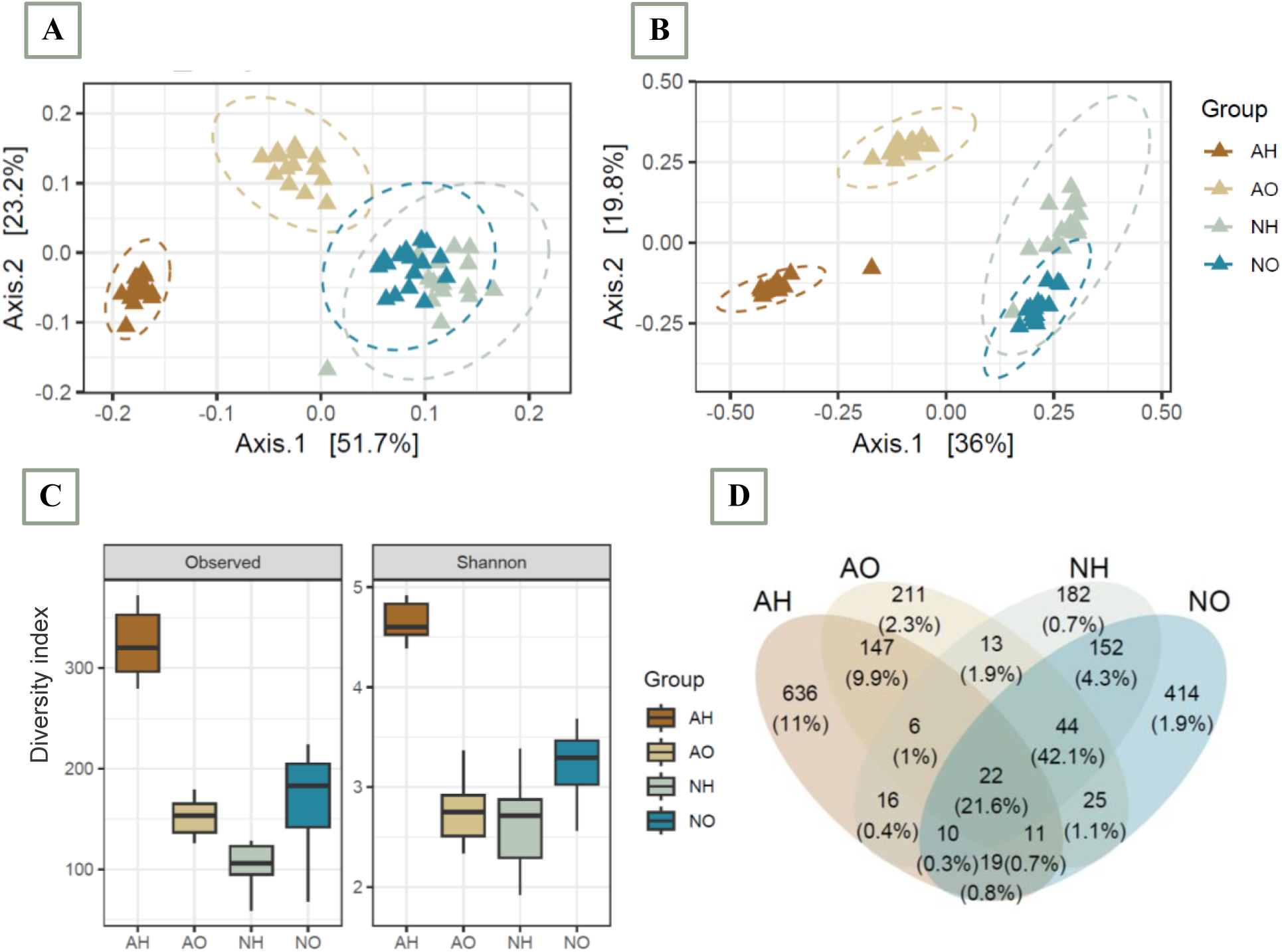
Root-associated bacterial communities are separable by different treatments presenting soil type (autoclaved vs. non-autoclaved) and temperature (high vs. optimum). A and B. PCoA plots using the Unifrac distance metrics (weighted and unweighted, respectively) indicate that the largest separation between bacterial communities of different treatments is soil type (PCo1) and the second-largest source of variation is the temperature (PCo2). C. Alpha-diversity measurements between different treatments indicate an increasing gradient in bacterial diversity of the rhizosphere in the autoclaved soil and high temperature. We estimated the Observed ASV richness and Shannon (H′) index in the bacterial communities across four different treatments shown with ± SE. D. Venn diagram showing the overlap between ASVs in response to different treatments based on the presence or absence of ASVs in 50% of the samples for each corresponding groups. Values are presented as the absolute number of ASVs and also as percentages (in parentheses).

Alpha diversity of the sequencing data and total ASV richness for different treatment conditions have been displayed in Figure 1C. Generally, alpha diversity indices including Shannon’s biodiversity and observed ASVs were significantly different among treatments (χ^2^ = 37.61, P = 3.47e-08, Shannon; χ^2^ = 43.33, P = 2.08e-09, observed; Kruskal-Wallis; Table S5). Although there were no statistically significant differences between genotypes (χ^2^ = 2.29, P = 0.51, Shannon; χ^2^ = 0.85, P = 0.83, observed, Kruskal-Wallis; Table S5, Fig. S1), a significant interaction effect of treatments and genotypes on rhizosphere bacterial diversity was observed (χ^2^ = 43.12, P = 0.0001, Shannon; χ^2^ = 45.8 P = 5.7e-05, observed, Kruskal-Wallis; Table S5).

A general view of the bacterial absolute numbers across different treatments showed that 21.6% of bacterial ASVs discovered in this analysis were common across different treatments (Fig. 1D). We observed a larger fraction of bacterial ASVs shared with the three treatments including NO, NH, and AO (42.1%). However, a large number of ASVs associated with the rhizobacterial communities presented in the AH treatment was depleted across other treatments (11%), indicating a larger separation between bacterial communities under autoclaved soil with high temperature. The relative abundance of various bacterial ASVs was differed between treatments (Fig. 2A). Although *Actinobacteria*, *Proteobacteria*, *Firmicutes*, and *Bacteroidota* were dominant in the rhizosphere community associated with all treatments, low-abundant phyla such as *Verrucomicrobia*, *Planctomycetota*, *Acidobacteriota*, *Chloroflexi*, *Myxococcota*, and *Gemmatimonadota* increased in the rhizosphere community of autoclaved soil with high temperature (Fig. 2A). The similarity of abundance was highest among the NH and AO. In contrast, AH and NO displayed the highest variation (Fig. 2A and S3). These observations suggest that a 10-degree gradient in air temperature (from 28 °C to 38 °C) may influence the rhizosphere bacterial communities of soybean like that of autoclaving.

**Figure 2.**
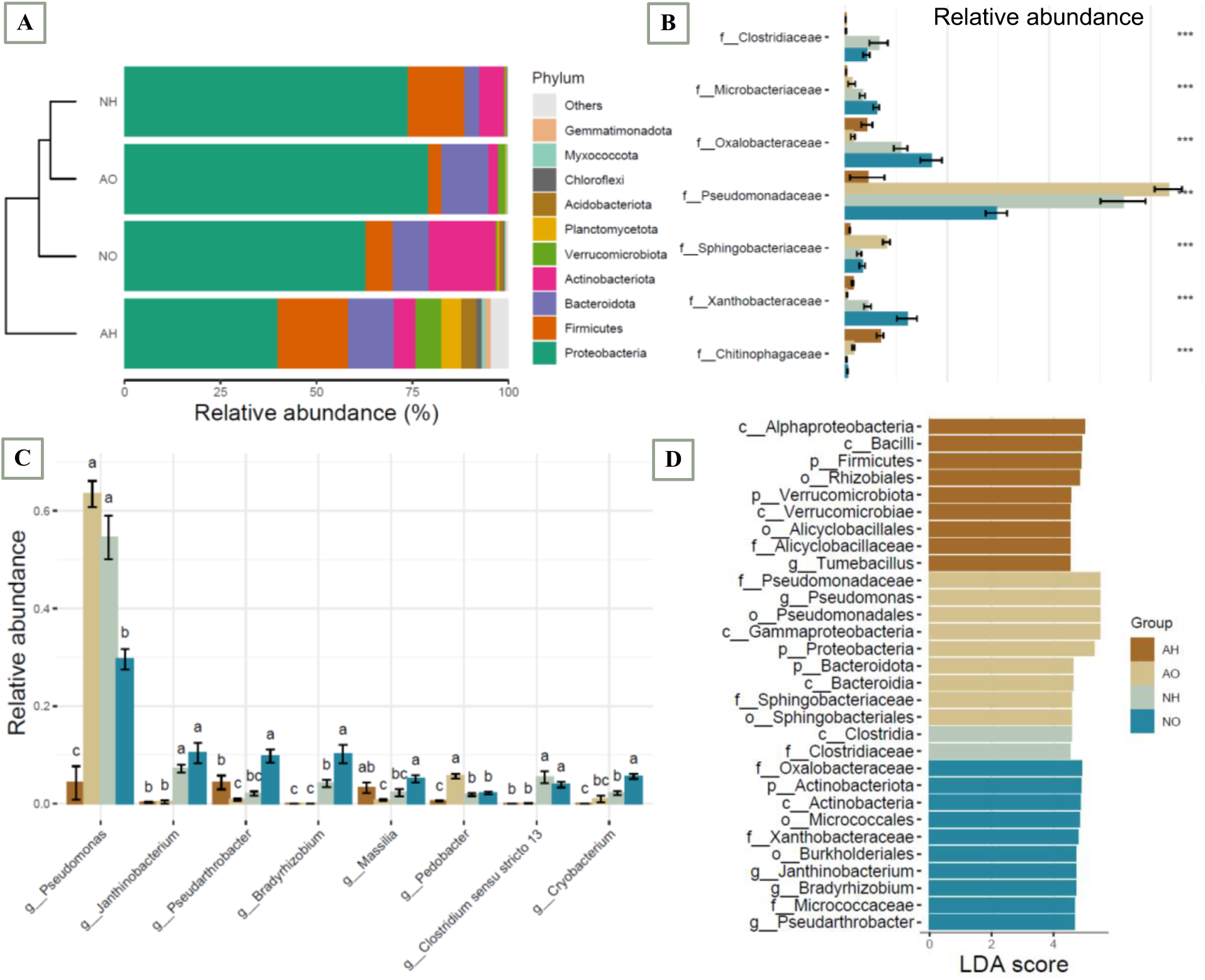
Rhizosphere bacterial composition responses to thermal stress and soil biotic disturbance.

Differential taxonomic analysis at the family level identified bacteria depleted in soybean rhizosphere under autoclaved soil with both ambient and high temperatures such as *Pseudomonadaceae*, *Sphingobacteriaceae*, *Micrococcaceae*, *Oxalobacteraceae*, *Xanthobacteraceae*, *Microbacteriaceae*, *Clostridiaceae*, and *Comamonadaceae*. However, *Sphingomonadaceae*, *Rhizobiaceae*, *Bacillaceae*, *Gemmatimonadaceae*, *Rhodobacteraceae*, *Bryobacteraceae*, *Chitinophagaceae*, and *Caulobacteraceae* were among families significantly enriched in AH (Fig. 2B and S4). At the genus level, the results revealed that several genera such as *Bradyrhizobium*, *Pseudarthrobacter*, *Pseudomonas*, *Cryobacterium*, *Colestridium sensu stricto 13*, *Masiilia*, and *Janthinobacterium* were highly differential among at least two different tretments (Fig. 2C). The differential abundant genera with an overall relative abundance larger than 0.05% for at least 50% of samples of each treatment are summarized in a relative abundance-prevalence plot depicted in Figure S5. We further identified biomarkers to differentiate between treatments using the linear discriminantanalysis effect size (LEfSe) algorithm (Fig. 2D). The results showed that the the family *Alicyclobacillaceae* and the genus *Tumebacillus* displayed higher relative abundances in the rhizosphere under AH. In contract, two families *Pseudomonadaceae* and *Sphingobacteriaceae* were among bacterial markers with high relative abundances in AO (Fig. 2B and C). The family *Clostridiaceae* displayed higher relative abundances in NH, wherease three families including *Oxalobacteraceae*, *Micrococcaceae*, *Xanthobacteraceae* and three genera *Janthinobacterium*, *Bradyrhizobium*, *Pseudarthrobacter* displayed higher relative abundances in the rhizosphere soil under NO.

A. Bar plot showing the most highly represented phyla in the rhizosphere of soybean plants across different treatments. Differential analysis results on bacteria at family and genus levels have been presented in bar plots B and C, respectively. D. The LEfSe analysis showed the biomarker taxa contributing to differences in the treatments with LDA log score threshold > 4.5 and P < 0.05.

### Fungi

A first comparative assessment of the effects of the grouping factors treatment and genotype on the fungal community diversity was made at the ASV level (Fig. 3A, B, and C). It showed that different treatments (R^2^ = 58.6%, p = 0.001, weighted; R^2^ = 40%, p = 0.001, unweighted; Table S4) and the interaction between treatment and plant genotype (R^2^ = 82.4%, p = 0.001, weighted; R^2^= 73%, p = 0.001, unweighted; Table S4) were primarily associated with dissimilarities between samples. Among two components of treatment including soil type and temperature, the fungal community composition responded most strongly to soil type (R^2^ = 39.4%, p = 0.001, weighted; R^2^ = 16.6%, p = 0.001, unweighted; PERMANOVA) followed by temperature (R^2^ = 4.2%, p = 0.22, weighted; R^2^ = 10.8%, p = 0.001, unweighted; PERMANOVA). These responses were clearly reflected in the PCoA plots, where samples were well separated along the first two axes according to these grouping factors (Fig. 3A and B). However, the differentiation of samples according to factor genotype was weak in the fungal dataset like that of bacterial communities (R^2^ = 5.3%, p = 0.95, weighted; R^2^ = 9.4%, p = 0.471, unweighted; Table S4). Analysis of the effects of treatment and genotype further revealed that fungal alpha diversity was responsive to treatment and the interaction between treatment and genotype similar to that of bacterial communities (Table S5). The strongest differences in estimated ASV richness and Shannon diversity for fungal communities were observed in relation to treatment (χ^2^ = 20.43, P = 0.0001, Shannon; χ^2^ = 17.25, P = 0.0006, observed; Kruskal-Wallis; Fig. 3C, Table S5) followed by the treatment-by-genotype interaction (χ^2^ = 25.38, P = 0.03, Shannon; χ^2^ = 22.24, P = 0.07, observed; Kruskal-Wallis; Table S5). Likewise, the grouping factors affected diversity more stronger in bacterial communities compared to fungal lineages (Table S5). We didn not found a difference in alpha diversity of fungal communities associated with the rhizosphere samples of different genotypes used in this study (Table S5 and Fig. S6).

**Figure 3.**
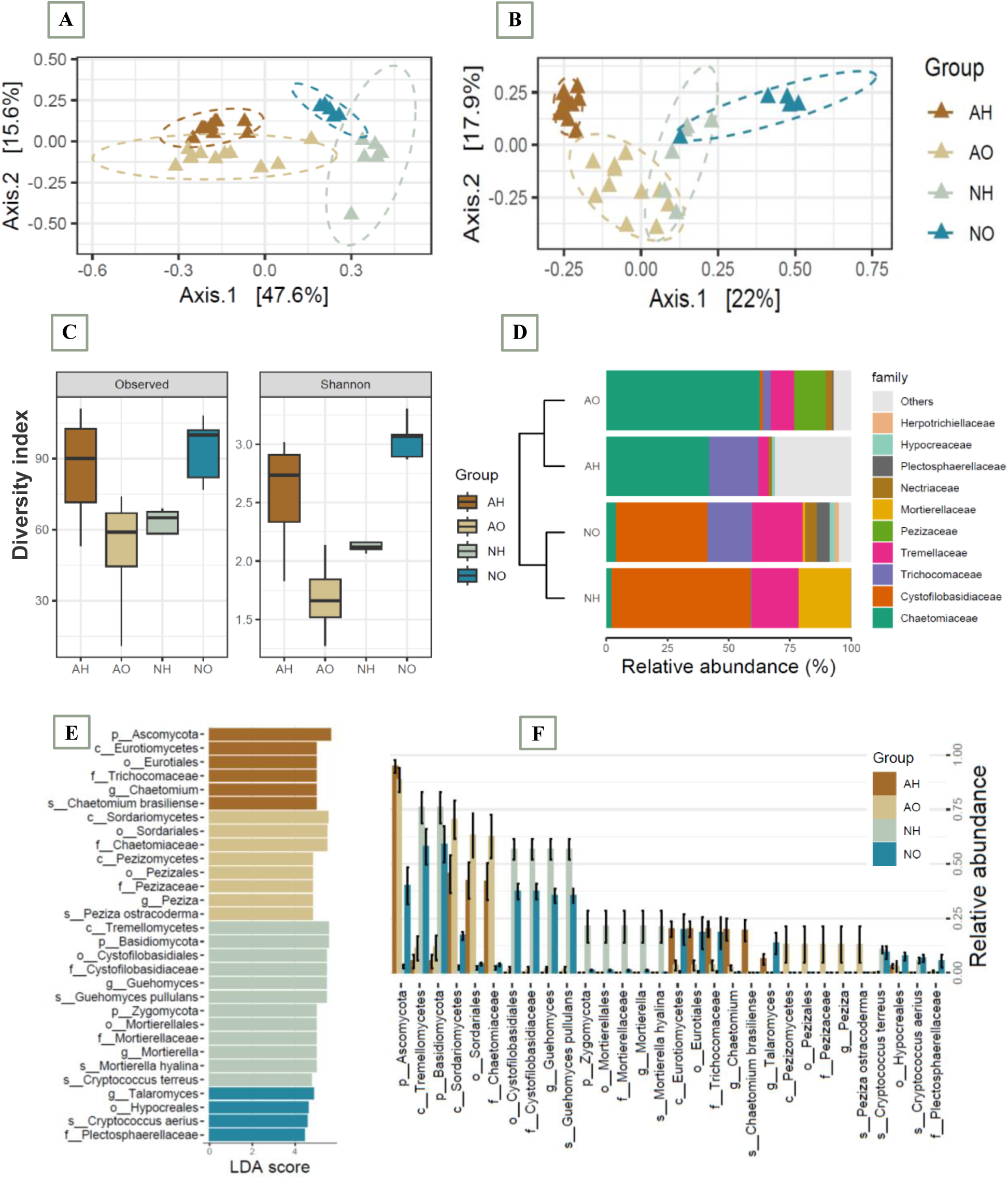
Dynamics of diversity, distribution patterns, and composition of soybean-associated fungal communities. PCoA plots using the Unifrac distance metrics including A. weighted and B. unweighted indicate that the rhizosphere-associated fungal communities are separable by different treatments. C. Alpha-diversity measurements between different treatments also show an significant differences in fungal diversity of the rhizosphere. The estimated observed ASV richness and Shannon (H′) index in the fungal communities across four different treatments has been shown with ± SE. D. Differential taxonomic analysis presenting relative abundance of each taxa at the family level showed differences in distribution patterns between treatments and confirmed that fungal communities of soybean rhizosphere clustered mainly based on soil biotic disturbance. E. The LEfSe analysis showed the biomarker fungi contributing to differences in the treatments with LDA log score threshold > 4.5 and P < 0.05. F. The abundance of biomarkers detected by LEfSe was visualized in the bar plot to compare different treatments.

The composition of the fungal communities in the soybean rhizosphere was dominated by *Ascomycota* (71.9%), *Basidiomycota* (25%), *Zygomycota* (3.1%) that are commonly present in soils. With regard to the experimental treatments a significantly higher relative abundance of *Ascomycota* in autoclaved compared to the non-autoclaved treatments was evident. In contrast, phyla *Basidiomycota* and *Zygomycota* were significantly enriched in the non-autoclaved treatments (Fig. S7). The compositional shift at the family level was also characterized by a prominent increase of a diverse group of fungi such as *Chaetomiaceae* and *Trichosporonaceae* and a decrease of *Cystofilobasidiaceae*, *Tremellaceae*, *Mortierellaceae*, *Nectriaceae*, *Plectosphaerellaceae*, and *Herpotrichiellaceae* in treatments with autoclaved soil compared to non-autoclaved soil in both termal levels, as presented in Figure 3D. Other fungal families such as *Hypocreaceae*, *Pezizaceae*, *Pleosporaceae*, *Pezizomycotinaceae*, *Phaeosphaeriaceae*, *Bionectriaceae*, *Leucosporidiaceae*, *rthrodermataceae*, *Hypocreaceae*, *Lasiosphaeriaceae*, *Helotiaceae*, *Sporidiobolaceae*, *Pseudeurotiaceae*, *Clavicipitaceae*, and *Trichosphaeriaceae* showed also differentially family-level relative abundance across different treatments (Fig. S8). Fungal biomarkers were further discovered using the LEfSe analysis (Fig. 3E and F). We found that the families *Trichosphaeriaceae* and *Chaetomiaceae* were strongly enriched in the soybean rhizosphere under AH. While, two families *Chaetomiaceae* and *Pezizomycotinaceae* were the most enriched biomarker taxa in AO. Furthermore, the abundance of families *Cystofilobasidiaceae* and *Mortierellaceae* were higher in the rhizosphere under NH compared to other treatments. In contrast, the family *Plectosphaerellaceae* was enriched in the rhizosphere of soybean genotypes in non autoclaved soil with ambient temperature (NO).

To understand the functional differences of microbial communities, we predicted the potential metabolic capacities of rhizosphere bacterial and fungal communities. The results displayed divergent metabolic capacities between soybean rhizosphere microbiome across different treatments (Fig. S9A and B). In bacteria, metabolic pathways related to biodegradation of xenobiotics, fatty acids, and benzoates, as well as those involved in metabolism of propanoate, phenylalanine, fatty acid and lipid were significantly enriched in the soybean rhizosphere of NO. However, pathways involved in nutrient transformation and transport, such as membrane and ABC transporters along with cellular processes and nucleotide metabolism were enriched in soybean rhizosphere under AH conditions. In contrast, bacterial functions related to plant-microbe interactions showed an increased accumulation in the rhizosphere of AO, such as signal transduction, two-component systems, cell motility along with biosynthesis of amino acids and secondary metabolites. Metabolic pathways involved in beta-alanine, amino acid, terpenoid, and polyketides were further enriched in NH treatment. In addition, the metabolic pathways for biofilm formation were accumulated in both AO and NH (Fig. S9A).

The application of FUNGuild showed that the total fungal ASVs associated with the key trophic modes pathotroph, saprotroph and symbiotroph, and their combinations (Fig. S9B). Totals of 17 functional guilds were identified, most of them affiliated with undefined saprotroph, symbiotroph, and endophyte forms (more than 75% of relative abundance). Plant pathogenic fungi also included almost 5% of relative abundance and revealed significant distributions across treatments. As NO showed the most fungal pathogenic forms, following by AO and NH, and the AH had the least relative abundance of pathogenic fungi. The other predicted functional guilds such as dung saprotroph, symbiotroph, endophyte, wood saprotroph, plant saprotroph revealed statistically significant differences between the treatments, and enriched more in autoclaved soil (Fig. S9B).

### Treatment-induced changes in root metabolites

We identified a total of 739 biochemicals, with 488 compounds of known identity and 251 compounds of unknown structural identity. Our analysis revealed significant metabolomic alterations in response to high temperature stress and soil biotic disturbances. Following log transformation and imputation of missing values, with the minimum observed value for each compound, ANOVA contrasts were used to identify biochemicals that differed significantly between experimental groups. Specifically, we observed 372 differentially accumulated metabolites (DAMs), highlighting the substantial biochemical shifts in soybean under these growing conditions (P < 0.05) (Fig. 4A). Metabolites such as flavonoids, organic acids and derivatives, amino acids and derivatives, nucleotides and derivatives, vitamins and derivatives, lipids, alkaloids, phenylpropanoids, terpenes, polyphenols, phenolic amines, and benzenoids were the main differential metabolites across different treatments (Table S6).

**Figure 4.**
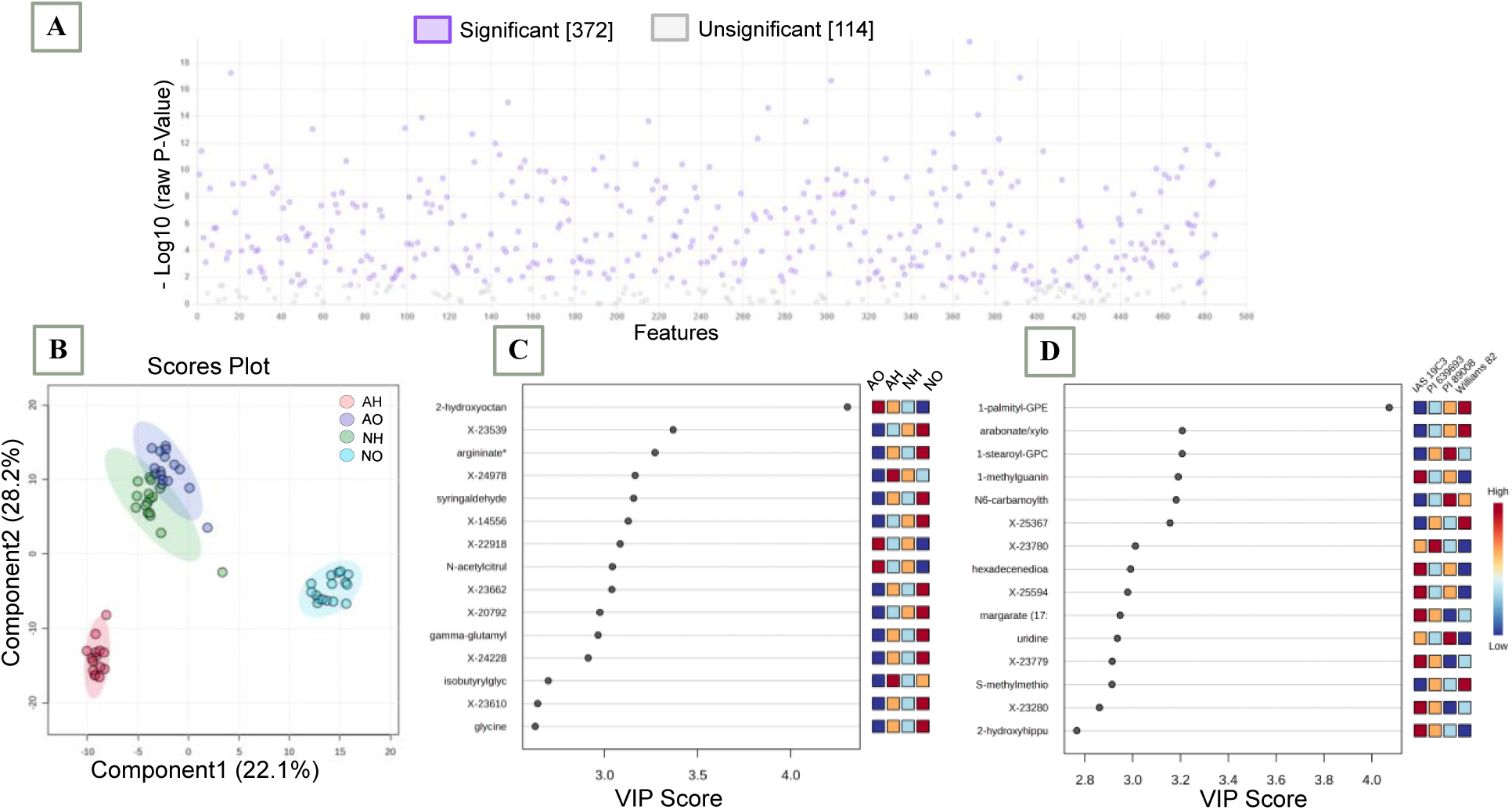
Plant metabolomic alterations in response to high temperature stress and soil biotic disturbances. A. Differentially accumulated metabolites (DAMs) with respect to heat stress and soil biotic disturbance. B. Partial least squares-discriminant analysis (PLS-DA) using metabolomic dataset at the 95% confidence interval. Variable importance in projection (VIP) score plot for the top 10 most important metabolite features contributing the separation identified by PLS-DA across C. different treatments, and D. genotypes. Red indicates higher abundance, while blue indicates lower abundance for a given metabolite.

The PLS-DA provided a clear separation of metabolite profiles between treatments, indicating that both heat stress and soil biotic disturbances induced distinct metabolic responses (Fig. 4B). The clusters mainly correlated to the thermal and microbial disturbance accounts, starting from the AH at one end and the NO at another end, which explained a high variation 22.1% of root metabolites between the two treatments in component 1. These treatments also showed a difference 28.2% with AO and NH treatments clustered more closely together in the middle of NH and AO. This suggests that the metabolome of soybean is highly sensitive to both heat stress and microbiome perturbations. We further performed the VIP score plot (2.0-5.0) to identify the top 10 metabolites responsible for the separation observed in the PLS-DA (Fig. 4C). Although, the amount of amino acids argininate, gamma-glutamylisoleucine, and glycine as well as phenylpropanoid syringaldehyde, enriched significantly in plant roots under NO treatment, metabolic profiles of AO showed a significant reduction of these metabolites, indicating the role of soil microbiome in plant metabolic modifications under ambient temperature. AH and NH also differed in contents of root metabolites. Under AH conditions, the fatty acid 2-hydroxyoctanoate, and N-acetylcitrulline, a derivative of amino acids, as well as a unkown metabolite increased significantly in the soybean roots, while NH showed an increase in the branched chain amino acid isobutyrylglycine, 2-hydroxyoctanoate, and several unknown metabolites. This observation indicates that the plant root metabolites probably have critical roles in soybean heat stress tolerance (Fig. 4C). Particularly, an enrichment of isobutryrylglycine and the unknown metabolite X24978 in NH and their meaningful decreased contents under AH. This evidence may indicate the plant microbiome contribution to alleviate thermal stress in soybean through metabolite reprogramming. A further analysis of the VIP showed that several metabolits belonged in fatty acids (margarate, hexadecenedioate), lipids (1-palmityl-GPE, 1-stearoyl-GPC), amino acids and amino sugars (S-methylmethionine, arabonate/xylonate), and the benzenoid 2-hydroxyhippurate as well as several substrates involved in nucleotide metabolism (1-methylguanine, N6-carbamoylthreonyladenosine, uridine) were the most important metabolites to explain the genotypic variation across different treatments (Fig. 4D).

### Relationships between plant metabolites and root microbiome

Since significant inter-omic correlations were found between soybean root microbiome and metabolome, pearson’s rank correlation analysis was performed between VIP metabolites and the root microbiome composition in both bacterial and fungal datasets (Fig. 5A and B). To identify relationships between bacterial and fungal communities and root metabolites, we used Pearson’s rank correlations and hierarchical clustering of differentially abundant rhizosphere ASVs and metabolites observed in the VIP score plots. In this analysis, we grouped both genotype and treatment-responsive rhizosphere metabolites and microbial ASVs by their degree of correlation, to identify clusters with similar behavior. Hierarchical clustering of the most responsive ASVs and differentially abundant metabolites revealed three large bacterial-metabolite clusters (Fig. 5A). The first cluster (at the top) contained microbial ASVs (n=7) and rhizosphere metabolites (n=11) that showed differential VIP score between tratments, including bacterial genera *Janthinobacter*, *Bradyrhizobium*, *Cryobacterium*, *Duganella*, *Flavobacterium*, *Massilia*, and *Pesudarthrobacter* and metabolites with N-rich compounds such as amino acids (glycine, argininate, gamma-glutamylisoleucine, and isobutyrylglycine), as well as syringaldehyde, and 2-hydroxyoctanoate. In addition, several genotype-responsive compounds such as hexadecenedioate, arabonate/xylonate, S-methylmethionine, 1-methylguanine, and 2-hydroxyhippurate showed a positive significant correlation with the genera from cluster 1. The majority of rhizosphere metabolites (54%) that co-varied with the ASVs from the first cluster were amino acid. This observation indicates that the ability to synthesize amino acids, in particular, can be linked to the differences between treatments influencing microbial colonization in the rhizosphere. Cluster 2 correlated the significant enrichment of two metabolites N6-carbamoylthreonyladenosine and uridine involved in nucleotide metabolism and the related bacterial ASVs (n=15). ASVs in this cluster were distinct from the ASVs identified in the Cluster 1 and include diverse microbial genera such as *Brevibacillus*, *Ammoniphilus*, *Ramlibacter*, *Chthoniobacter*, *Fimbriiglobus*, *Flavisolibacter*, *Sphingoaurantiacus*, *Gemmatimonas*, *Microvirg*a, *Tabrizicola*, *Tumebacillus*, *Bryobacter*, *Nibrella*, *Pedomicrobiome*, and *Prosthecomicrobiome*. However, cluster 3 presented ASVs with a negative correlation with plant metabolites. Specially two lineages including *Limnohabitans* and *Pseudomonas* associated with a significant reduction in root metabolites such as amino acids (glycine, gamma-glutamylisoleucine, N-acetylcitrulline, and isobutyrylglycine), and metabolites involved in nucleotide metabolism (N6-carbamoylthreonyladenosine, 1-methylguanine, and uridine).

**Figure 5.**
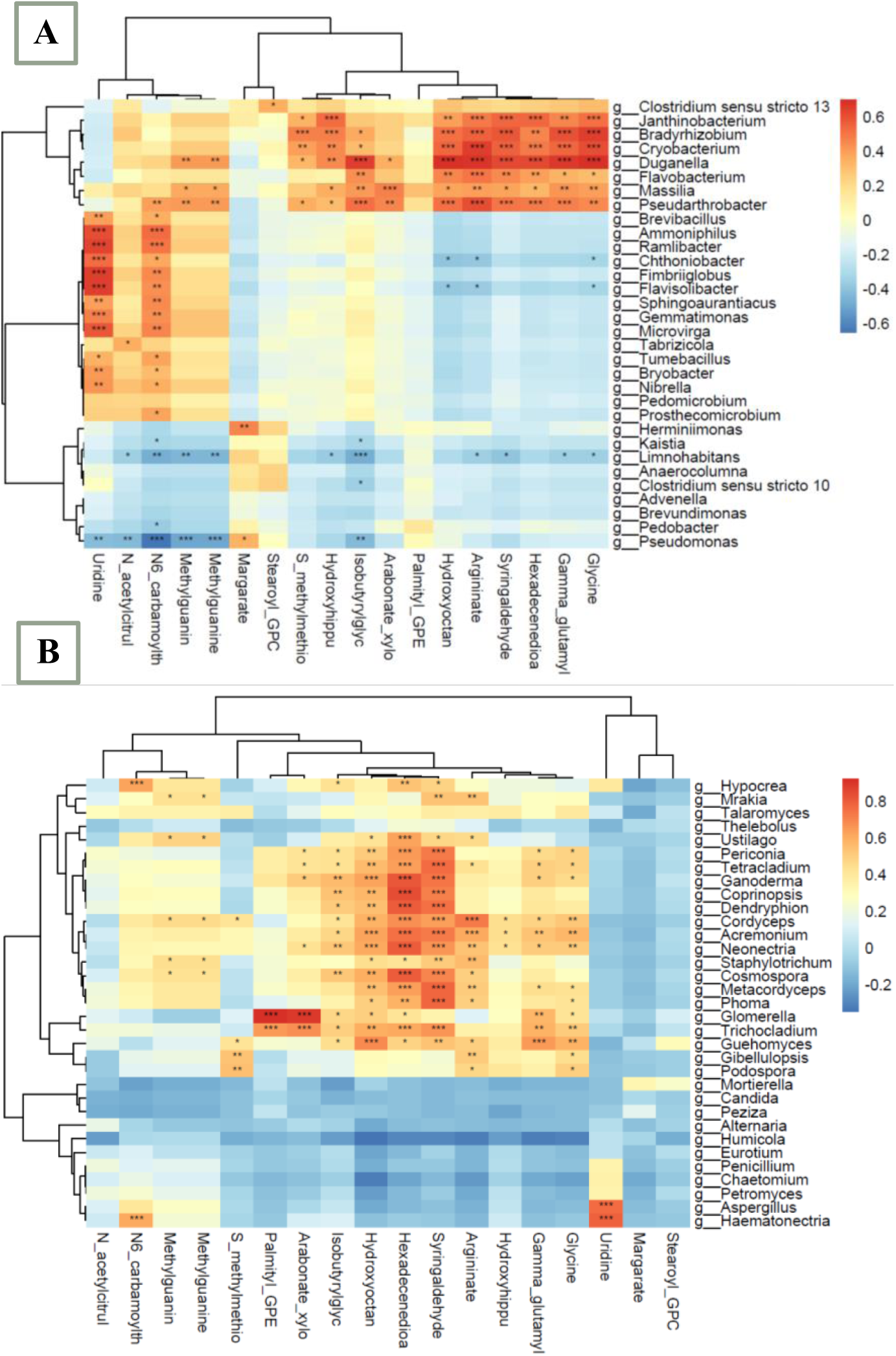
Heat map of the correlation analysis between top 10 soybean root metabolites differentially accumulated across different treatments and genotypes and A. bacterial and B. fungal community compositions. Each grid represents the correlation between the two attributes, and different colors represent the size of the correlation coefficient between the attributes. *p < 0.05; **p < 0.01.

The correlation analysis also showed that root metabolite was significantly correlated with fungal composition in the rhizosphere (Fig. 5B). ASVs and differentially abundant metabolites revealed two large fungal-metabolite clusters. Cluster 1 included treatment-responsive metabolites and microbial ASVs (n=22). Fungal genera such as *Periconia*, *Tetracladium*, *Ganoderma*, *Coprinopsis*, *Dendryphion*, *Cordyceps*, *Acremonium*, *Neonectria*, *Staphylotrichum*, *Cosmospora*, *Metacordyceps*, *Phoma*, *Trichocladium*, and *Guehomyces* were among ASVs presented in this cluster. In addition to the treatment-responsive metabolites, there was a positive correlation between some genotype-specific metabolites such as hexadecenedioate, 1-palmityl-GPE, S-methylmethionine, arabonate/xylonate, 2-hydroxyhippurate, and 1-methylguanine with ASVs of cluster 1. Cluster 2 contained ASVs including *Aspergillus* and *Haematonectria* and root metabolites N6-carbamoylthreonyladenosine, and uridine that were among top-10 metabolites correlated to soybean genotypes.

Linear regression model was then used to determine if the particular metabolites are significantly different in abundance between treatments. The results showed that among all VIP-metabolites responsive to heat stress and soil microbial disturbance: 2-hydroxyoctanoate, isobutyrylglycine, argininate, and syringaldehyde are the best predictor of metabolic features correlated to microbiome dissimilarity across treatments (Fig. S10). For soybean genotypes, only average amount of metabolites N6-carbamoylthreonyladenosine, 1-methylguanine, uridine and margarate were weakly, yet significantly correlated to rhizosphere microbiome in Williams82 and PI639693 (Fig. S11). Mantel test results further confirmed a significant correlation between features involved in nucleotide metabolism and bacterial community (N6-carbamoylthreonyladenosine, P = 0.009; 1-methylguanine, P = 0.04; uridine, P = 0.009). Similar results were found in genotype-responsive metabolites correlated to fungal communities and the metabolites involved in nucleotide metabolism including N6-carbamoylthreonyladenosine and uridine were weakly and negatively correlated to plant microbiome in Williams82 (Fig. S12). However no significant metabolite was observed for the VIP metabolites including frequency and intensity of rhizosphere colonization by fungi across different treatments.

### Relationships between root phenotypic traits and plant microbiome

The differential significance analysis of nodule and root anatomical traits including percent pitch, nodulation efficiency, fresh shoot weight, nodule weight, and the number of nodules significantly varied between rhizosphere bacterial and fungal communities of soybean and heavily influenced by treatments (Fig. 6, and Fig. S13). The nodule number, weight, and efficiency was highly compromised by the high temperature stress and microbiome disturbance. However, the amounts of xylem area, pitch area, and total area revealed no significant differences across treatments. The optimum temperature positively influenced shoot fresh weight, number of nodules, nodule weight, and nodule efficiency in both soil types (Fig. 6). Redundancy analysis (RDA) based on the specific relationships and the bacterial community structure further confirmed the differences in root phenotypic traits among treatments (Fig. 7). The bacterial community composition of AH was correlated with percent of pitch, pitch area, xylem area, and total area of roots. In contrast, rhizobacterial communities of NO were well separated from AO associated with a relatively higher shoot fresh weight. In addition, nodulation efficiency, number of nodules, and nodule weight had a more positive relationship with bacterial community composition of NO than AO. Our results also showed that various members of the bacterial communities such as *Bradyrhizobium*, *Janthinobacterium*, *Cryobacterium*, *Pseudarthrobacter*, and *Clostridium sensu stricta* were among the genera positively correlated with nodulation efficiency, number of nodules, and nodule weight. However, the aboundance of *Pedobacter* and *Pseudomonas* was positively associated with fresh shoot weight, and *Flavisolibacter*, *Tumebacillus*, and *Microvirga* were among members correlated to pitch area and percent of pitch (Fig. 7A and Fig. S14A).

**Figure 6.**
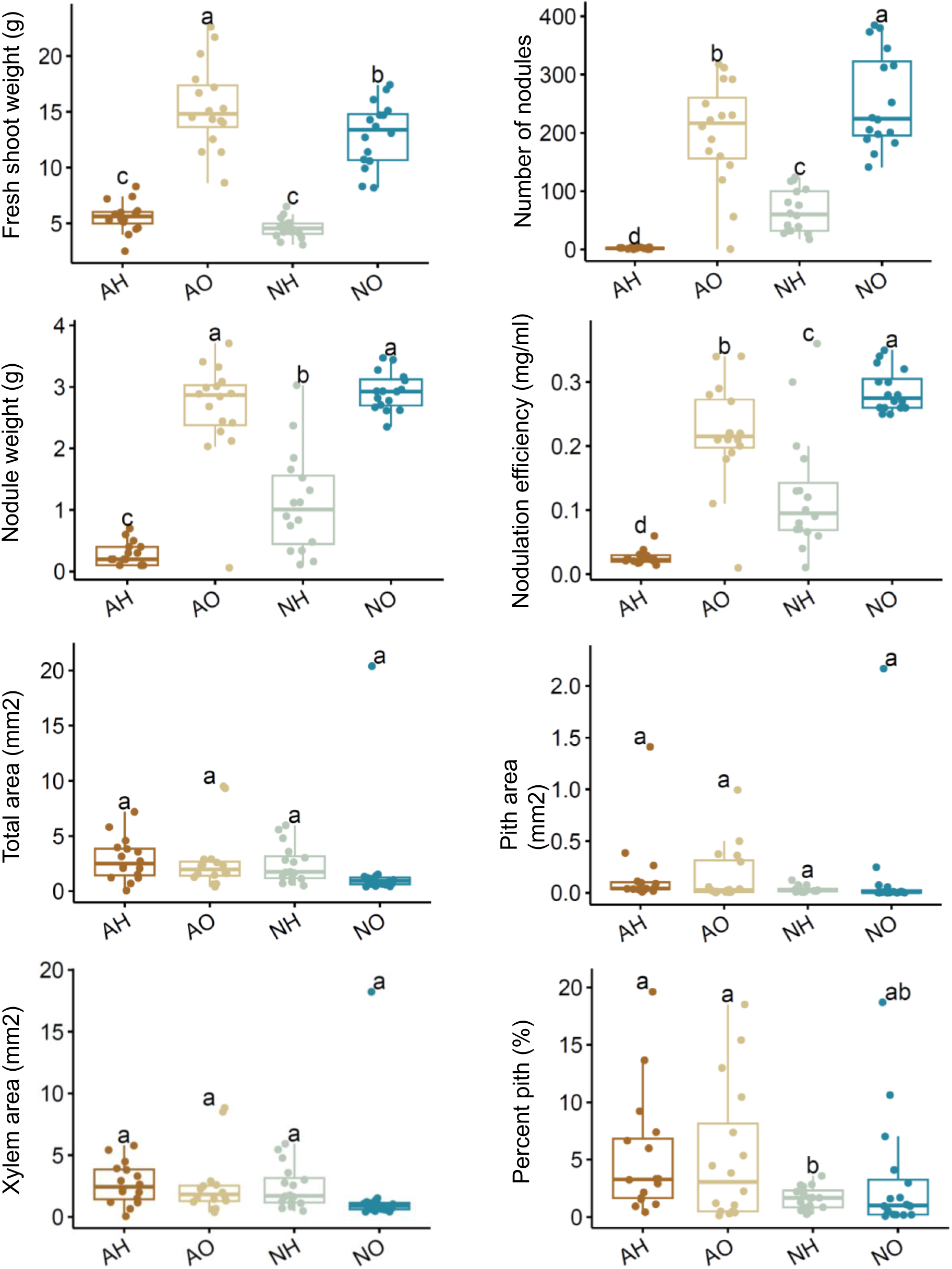
The differential significance analysis of nodule and anatomical variables in the rhizosphere bacterial communities of soybean submitted to different treatments.

**Figure 7.**
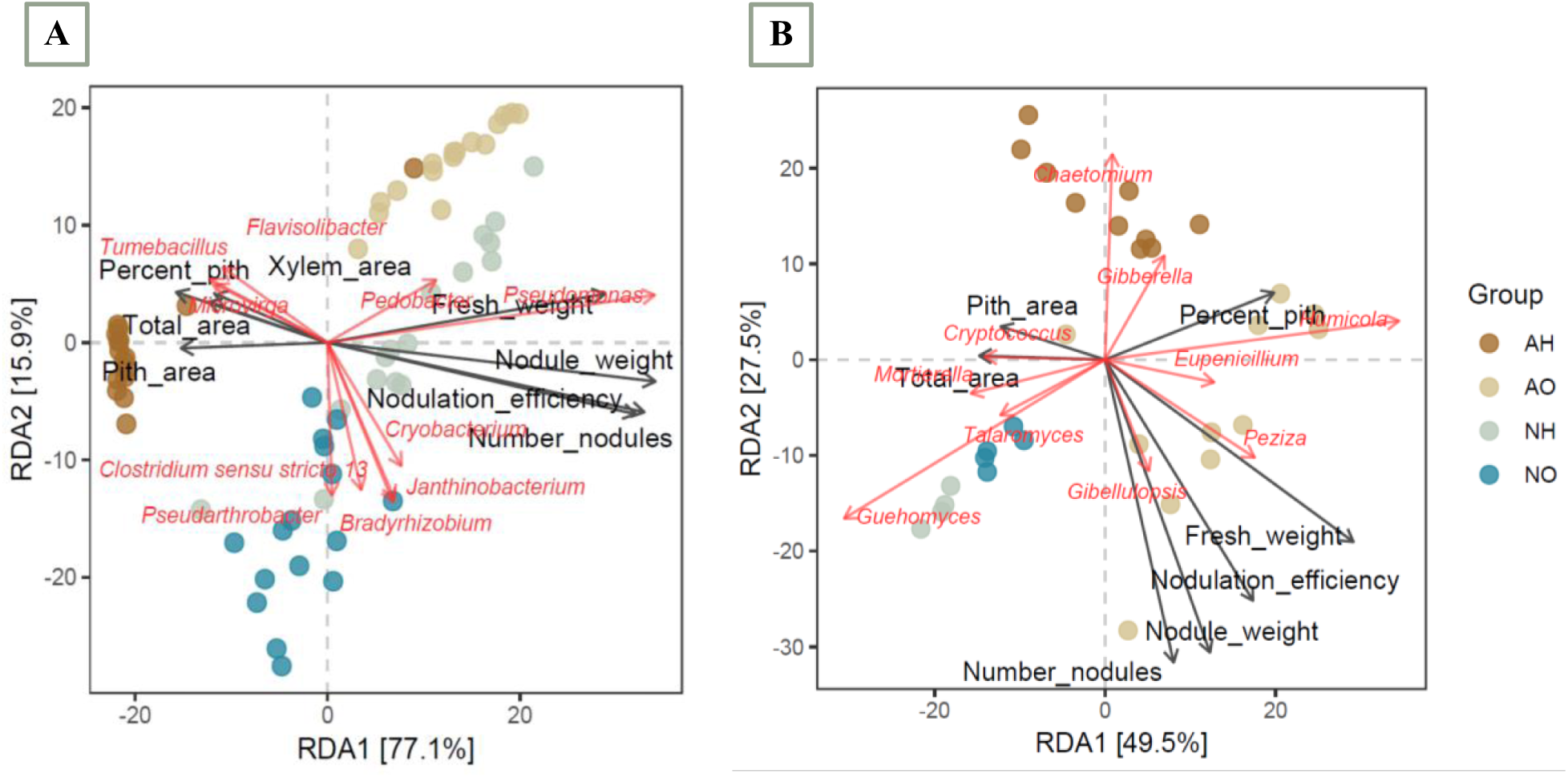
Redundancy analysis (RDA) to show the correlation between the bacterial community structure and plant phenotypic traits. RDA of A. bacterial and B. fungal communities generated using the Bray-Curtis distance, across different treatments, abundant genus (red arrows), and plant phenotypic factors (black arrows) indicate the dominant communities and plant phenotypic factors, respectively.

The relationships between plant phenotypic variables and fungal compositions were further investigated. RDA displayed a marked correlation between plant attributes and fungal compositions (Fig. 7B). The first two axes of the RDA accounted for 49.5% and 27.5%, respectively, of the total variation in the data. Like bacterial communities, the fungal compositions of AO and NO treatments were different from those of the AH treatment, and this was correlated to a relatively high shoot fresh weight, number of nodule, nodule weight, nodulation efficiency, and percent of pitch under optimum temperature (Fig. 7B). In contrast, the fungal compositions of AH were completely separated from that of the AO and NO treatments, which had a relatively high correlation with pitch area. Further, various members of the fungal genera such as *Eupenecillium*, *Peziza*, *Gibellulopsis*, *Talaromyces*, and *Guehomyces* positively correlated with shoot fresh weight and nodule variables in AO and NO treatments. While, a strong correlation between fungal genus *Periconia*, *Fusarium*, *Ustilago*, and *Neonectria* and plant phenotypic traits including xylem area, total area, and percent of pitch was also detected (Fig. S14B).

## Discussion

Understanding the complex interplay between plant roots, soil microbial communities, and environmental stressors is critical to improve the soil and plant health [59]. This study presents the integrative assessments on how heat stress and soil microbial disturbance influence rhizosphere microbial assembly, root metabolomic reprogramming, and nodulation in soybean. Through amplicon sequencing, non-targeted metabolomics, and root phenotyping, we provide evidence that microbial communities are critical modulators of plant resilience, with cascading effects on belowground symbiosis and metabolic architecture.

### Soil microbiome integrity buffers soybean response to heat stress

Our results reveal that the presence of a native, undisturbed soil microbiome significantly mitigates the adverse effects of high temperature on rhizosphere diversity and nodulation efficiency. Autoclaving-induced microbial disturbance led to a collapse in microbial richness and community stability, as evidenced by both alpha and beta diversity metrics, particularly under high-temperature stress (AH treatment). PERMANOVA analysis showed that treatment conditions explained the majority of variation in microbial community structure, while genotype contributed only modestly. This finding suggests that environmental factors, particularly microbial presence and thermal conditions, are the dominant drivers of rhizosphere assembly, aligning with emerging paradigms in holobiont theory that emphasize the environment’s role in shaping host-associated microbiomes [60].

The core symbiotic genera such as *Bradyrhizobium* and *Janthinobacterium* were depleted under microbial disturbance and heat stress conditions. These taxa are not only pivotal for biological nitrogen fixation but are also associated with microbial consortia that regulate host immunity and stress signaling [61]. Their reduced abundance likely contributed to the sharp decline in nodulation metrics and leghemoglobin content under AH conditions. In contrast, the NH and NO treatments, which maintained intact microbial communities, exhibited enhanced nodulation, suggesting that microbial buffering capacity plays a central role in sustaining symbiotic functions during heat exposure.

### Metabolomic reprogramming as a response to microbial and thermal cues

Root metabolite profiles revealed pronounced reprogramming under the combined influence of temperature and soil microbial status. A total of 372 differentially accumulated metabolites (DAMs) were identified across treatments, representing a wide array of biochemical classes including amino acids, organic acids, phenylpropanoids, lipids, and nucleotides. PLS-DA and VIP metrics indicated that specific metabolites such as isobutyrylglycine, syringaldehyde, and 2-hydroxyoctanoate were strongly enriched in treatments with intact microbiomes, particularly under NH conditions.

The accumulation of amino acid-derived compounds and signaling metabolites under non-autoclaved conditions suggests that the presence of beneficial microbes supports stress-adaptive metabolite synthesis. In contrast, autoclaved soils lacking native microbial partners showed depletion of key metabolites involved in redox balance, osmoprotection, and signaling. Notably, the observed decline in nucleotide-related metabolites (uridine and N6-carbamoylthreonyladenosine) under microbial disturbance points to potential disruptions in transcriptional and translational regulation under stress. These findings reinforce the concept of the rhizosphere as a metabolically active interface, where microbial presence enhances plant biochemical plasticity.

### Microbe-metabolite interactions shape rhizosphere dynamics

By integrating microbial ASV profiles with metabolomic data, we identified tightly co-varying clusters that underpin the plant-microbe-metabolite nexus. Bacterial genera such as *Bradyrhizobium, Janthinobacterium, Cryobacterium,* and *Pseudarthrobacter* showed strong positive correlations with amino acids and lipid derivatives that were differentially enriched across treatments. These interactions suggest a possible two-way communication system, wherein plants exude specific metabolites to recruit beneficial microbes, which in turn modulate host stress responses [62]. Similarly, fungal taxa such as *Peziza, Guehomyces,* and *Talaromyces* were associated with metabolite profiles characteristic of high nodulation and biomass accumulation. These fungi may serve supportive or facilitative roles, such as modulating nutrient availability, secreting signaling molecules, or interacting synergistically with rhizobia. The clustering of ASVs and metabolites into functionally distinct modules highlights the potential for developing targeted microbial consortia that can enhance plant resilience through biochemical complementation.

### Functional restructuring of rhizosphere microbiota under stress

Functional predictions using Tax4Fun2 and FUNGuild revealed that microbial disturbance not only alters community composition but also drives functional reorganization within the rhizosphere. In autoclaved soils, we observed an enrichment of microbial pathways associated with stress response, nutrient limitation, and opportunistic colonization. These included ABC transporters, DNA repair mechanisms, and simple carbon metabolism. Conversely, in non-autoclaved soils, pathways involved in amino acid biosynthesis, biofilm formation, and two component regulatory systems were upregulated functions typically associated with symbiosis and microbial cooperation. The functional shift in fungal trophic modes was also notable. Saprotrophic and pathotrophic fungi became more abundant in autoclaved soils, suggesting that microbial disturbance creates ecological niches for opportunistic taxa, potentially exacerbating stress responses in host plants. In contrast, symbiotrophic fungi were enriched in non-autoclaved treatments, underscoring the ecological importance of an intact microbiome in maintaining functional redundancy and resilience in the rhizosphere.

### Microbiome-driven modulation of root traits and symbiosis

The physiological implications of microbial shifts were evident in root phenotyping data. RDA revealed that microbial community composition was tightly linked to root phenes such as nodule number, leghemoglobin content, and biomass. Notably, the AH treatment characterized by high temperature and microbial depletion showed the weakest nodulation response and poorest shoot growth, despite relatively stable root anatomy. This suggests that functional, rather than structural, attributes of the root system are more sensitive to microbial and thermal perturbation. Interestingly, while root xylem and pith areas remained largely unaltered, the percent pith area and nodulation-related traits showed strong correlations with key microbial taxa. This indicates that microbial-mediated processes influence specific root compartments involved in water transport, nutrient exchange, and hormonal signaling. These relationships point toward a nuanced interplay where microbes modulate specific root developmental trajectories, potentially through hormonal or metabolite-mediated pathways.

## Conclusion

Our study demonstrates that soil microbiome plays a pivotal role in shaping soybean responses to heat stress by modulating microbial diversity, root metabolite profiles, and symbiotic functions. The disruption of native microbiota impairs plant metabolic plasticity and nodulation, thereby compromising resilience. These insights underscore the necessity of incorporating microbiome conservation and engineering into crop breeding and soil management frameworks. As we continue to face increasingly unpredictable environmental stressors, such holistic, systems level approaches will be essential for continuous global food production and ecosystem stability. The findings from this study may have implications for sustainable agriculture and crop improvement strategies, preserving native soil microbiota is not merely beneficial but essential for optimal plant performance under environmental stress.

