## Supplementary files for "Heat Stress and Soil Microbial Disturbance Influence Soybean Root Metabolite, Microbiome Profiles, and Nodulation"

### Supplementary Information

#### A- Supplementary Tables

**Table S1**. Physical properties of the autoclaved and non-autoclaved soils (Soil texture class: Loamy)

| Soil Type | Clay | Silt | Sand | Very fine sand | Fine sand | Medium sand | Coarse sand | Very coarse sand |
|---|---|---|---|---|---|---|---|---|
| | µm | | | | | | | |
| Autoclaved | 22.05 | 31.44 | 46.35 | 14.25 | 12.05 | 12.20 | 6.81 | 1.05 |
| Non-autoclaved | 22.11 | 32.46 | 45.31 | 14.66 | 11.81 | 11.64 | 6.31 | 0.89 |

**Table S2**. Chemical properties of the autoclaved and non-autoclaved soils

| Soil type | Non-autoclaved | | | | Autoclaved | | | |
|---|---|---|---|---|---|---|---|---|
| Replication | 1 | 2 | 3 | 4 | 1 | 2 | 3 | 4 |
| Organic Matter (%) | 2 | 2.3 | 2.2 | 2.3 | 2.4 | 2.4 | 2.5 | 2.4 |
| Phosphorus P1 (Weak Bray) (ppm) | 16 | 19 | 17 | 17 | 31 | 36 | 36 | 31 |
| Phosphorus P2 (Strong Bray) (ppm) | 27 | 29 | 27 | 39 | 55 | 55 | 55 | 59 |
| Potassium (ppm) | 216 | 219 | 224 | 223 | 198 | 206 | 213 | 210 |
| Magnesium (ppm) | 257 | 287 | 283 | 288 | 283 | 289 | 300 | 299 |
| Calcium (ppm) | 1787 | 1872 | 1851 | 1925 | 1832 | 1887 | 1954 | 1945 |
| Sodium (ppm) | 9 | 12 | 13 | 12 | 13 | 14 | 13 | 15 |
| Soil pH | 6.1 | 6.1 | 6.1 | 6.1 | 5.7 | 5.8 | 5.7 | 5.8 |
| pH Buffer Index | 6.7 | 6.7 | 6.7 | 6.7 | 6.6 | 6.6 | 6.6 | 6.6 |
| Cation Exchange Capacity (meq/100g) | 13.6 | 14.4 | 14.2 | 14.7 | 15.3 | 15.3 | 16.3 | 15.8 |
| K (%) | 4.1 | 3.9 | 4 | 3.9 | 3.3 | 3.5 | 3.4 | 3.4 |
| Mg (%) | 15.7 | 16.6 | 16.6 | 16.3 | 15.4 | 15.7 | 15.3 | 15.8 |
| Ca (%) | 65.7 | 65 | 65.2 | 65.5 | 59.9 | 61.7 | 59.9 | 61.6 |
| H (%) | 14.2 | 14.1 | 13.8 | 13.9 | 21 | 18.7 | 21.1 | 18.8 |
| Na (%) | 0.3 | 0.4 | 0.4 | 0.4 | 0.4 | 0.4 | 0.3 | 0.4 |
| Nitrate-N (ppm) | 15 | 15 | 15 | 15 | 12 | 14 | 13 | 12 |
| Nitrate-N (lbs/A) | 27 | 27 | 27 | 27 | 22 | 25 | 23 | 22 |
| Sulfur (ppm) | 18 | 12 | 15 | 15 | 28 | 32 | 34 | 60 |
| Zinc (ppm) | 3.5 | 3.7 | 3.5 | 3.5 | 2.1 | 2.1 | 2.3 | 2.3 |
| Manganese (ppm) | 7 | 7 | 8 | 8 | 142 | 162 | 157 | 163 |
| Iron (ppm) | 38 | 37 | 40 | 36 | 22 | 21 | 21 | 24 |
| Copper (ppm) | 0.6 | 0.6 | 0.7 | 0.6 | 0.6 | 0.7 | 0.6 | 0.8 |
| Boron (ppm) | 0.2 | 0.2 | 0.2 | 0.2 | 0.2 | 0.2 | 0.2 | 0.2 |
| Soluble salts (mmhos/cm) | 0.2 | 0.2 | 0.2 | 0.2 | 0.4 | 0.5 | 0.4 | 0.4 |

**Table S3**. Changes in chemical properties of autocleved and non-autoclaved soils

| Soil chemical properties | Non-autoclaved | Autoclaved | P-value |
|---|---|---|---|
| | Mean-value | | |
| Organic Matter (%) | 2.2 | 2.425 | 0.04355 |
| Phosphorus P1 (Weak Bray) (ppm) | 17.25 | 33.5 | 0.0004376 |
| Phosphorus P2 (Strong Bray) (ppm) | 30.5 | 56 | 0.001517 |
| Potassium (ppm) | 220.5 | 206.75 | 0.01565 |
| Magnesium (ppm) | 278.75 | 292.75 | 0.1599 |
| Calcium (ppm) | 1858.75 | 1904.5 | 0.2989 |
| Sodium (ppm) | 11.5 | 13.75 | 0.07575 |
| Soil pH | 6.1 | 5.75 | 0.001208 |
| pH Buffer Index | 6.7 | 6.6 | 1 |
| Cation Exchange Capacity (meq/100g) | 14.225 | 15.675 | 0.004847 |
| K (%) | 3.975 | 3.4 | 0.0001111 |
| Mg (%) | 16.3 | 15.55 | 0.02954 |
| Ca (%) | 65.35 | 60.775 | 0.001627 |
| H (%) | 14 | 19.9 | 0.002677 |
| Na (%) | 0.375 | 0.375 | 1 |
| Nitrate-N (ppm) | 15 | 12.75 | 0.01822 |
| Nitrate-N (lbs/A) | 27 | 23 | 0.01094 |
| Sulfur (ppm) | 15 | 38.5 | 0.0462 |
| Zinc (ppm) | 3.55 | 2.2 | 2.54E-06 |
| Manganese (ppm) | 7.5 | 156 | 0.0000728 |
| Iron (ppm) | 37.75 | 22 | 0.00001 |
| Copper (ppm) | 0.625 | 0.675 | 0.4012 |
| Boron (ppm) | 0.2 | 0.2 | 1 |
| Soluble salts (mmhos/cm) | 0.2 | 0.425 | 0.002896 |

**Table S4**. The Permutational multivariate analysis of variance (PERMANOVA) to test the effect of soil type, temperature, and genotype, and their interactions on the bacterial communities associated with soybean. Numbers in sub-indices indicate the degrees of freedom and residuals of each F test.

| Dataset | Factor | Weighted | | | Unweighted | | |
|---|---|---|---|---|---|---|---|
| | | F | $R^2$ | P-value | F | $R^2$ | P-value |
| **Bacteria** | Treatment $_{3, 60}$<br>Genotype $_{3,60}$<br>Genotype x Treatment $_{15,48}$ | 66.028<br>0.3156<br>14.742 | 0.76752<br>0.01554<br>0.82164 | 0.001<br>0.984<br>0.001 | 34.881<br>0.4739<br>8.1081 | 0.6355<br>7<br>0.0231<br>5<br>0.7170<br>2 | 0.001<br>0.987<br>0.001 |
| **Fungi** | Treatment $_{3, 28}$<br>Genotype $_{3,28}$<br>Genotype x Treatment $_{15,16}$ | 13.211<br>0.5217<br>4.9999 | 0.58601<br>0.05293<br>0.82417 | 0.001<br>0.95<br>0.001 | 6.2471<br>0.9717<br>2.8919 | 0.4009<br>6<br>0.0943 | 0.001<br>0.471<br>0.001 |

| | | | | | | |
|---|---|---|---|---|---|---|
| | | | | | | 0.7305 4 |

**Table S5**. The effect of soil type, temperature, and plant genotype and their two-way interaction on the alpha-diversity of bacterial communities associated with the rhizosphere of soybean. Statistical support was done with the function "kruskall.test" in the R base. All P values were corrected for multiple comparisons using the FDR correction. Numbers in sub-indices indicate the degrees of freedom and residuals of each F test.

| Dataset | Factor | Shannon | | Observed | |
|---|---|---|---|---|---|
| | | Chi-square ($\chi 2$) | P-value | Chi-square ($\chi 2$) | P-value |
| **Bacteria** | Treatment $_{3, 60}$<br>Genotype $_{3,60}$<br>Genotype x Treatment$_{15,48}$ | 37.616<br>2.2999<br>43.125 | 3.407e-08<br>0.5125<br>0.0001 | 43.338<br>0.8549<br>45.807 | 2.086e-09<br>0.8363<br>5.706e-05 |
| **Fungi** | Treatment $_{3, 27}$<br>Genotype $_{3,27}$<br>Genotype x Treatment $_{14,16}$ | 20.433<br>0.088206<br>25.385 | 0.000138<br>0.9932<br>0.03095 | 17.25<br>2.1627<br>22.245 | 0.000627<br>0.5393<br>0.07372 |

B- Supplementary Figures

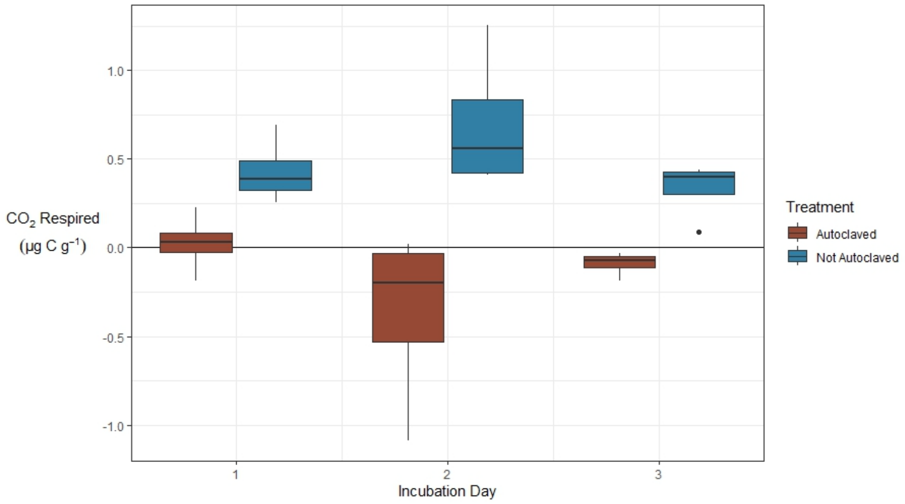

**Fig. S1**. **Microbial respiration rate for autoclaved and non-autoclaved soils** (values shown in mean ± SE; n = 4). Different lowercase letters indicate significant differences at $P < 0.05$ among the Soil type. *P < 0.05, **P < 0.01, ***P < 0.001. (in a box, we can show the results of the statistics for Soil type, Time and Soil type * Time).

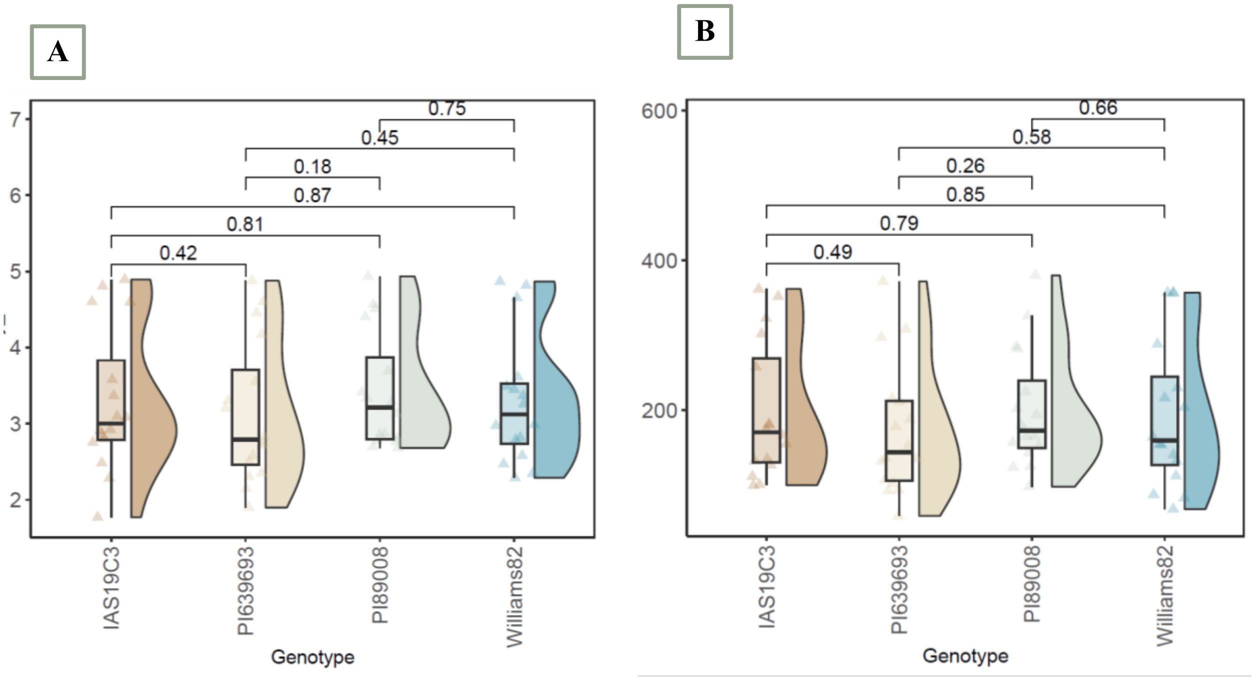

**Fig. S2**. **Alpha diversity measurement within the rhizosphere displayed small but not significant differences between the genotypes**. Estimated **A.** Shannon index and **B.** observed ASV richness in the bacterial communities associated with the four soybean genotypes across four different treatments shown with ± SE and pairwise wilcox comparisons.

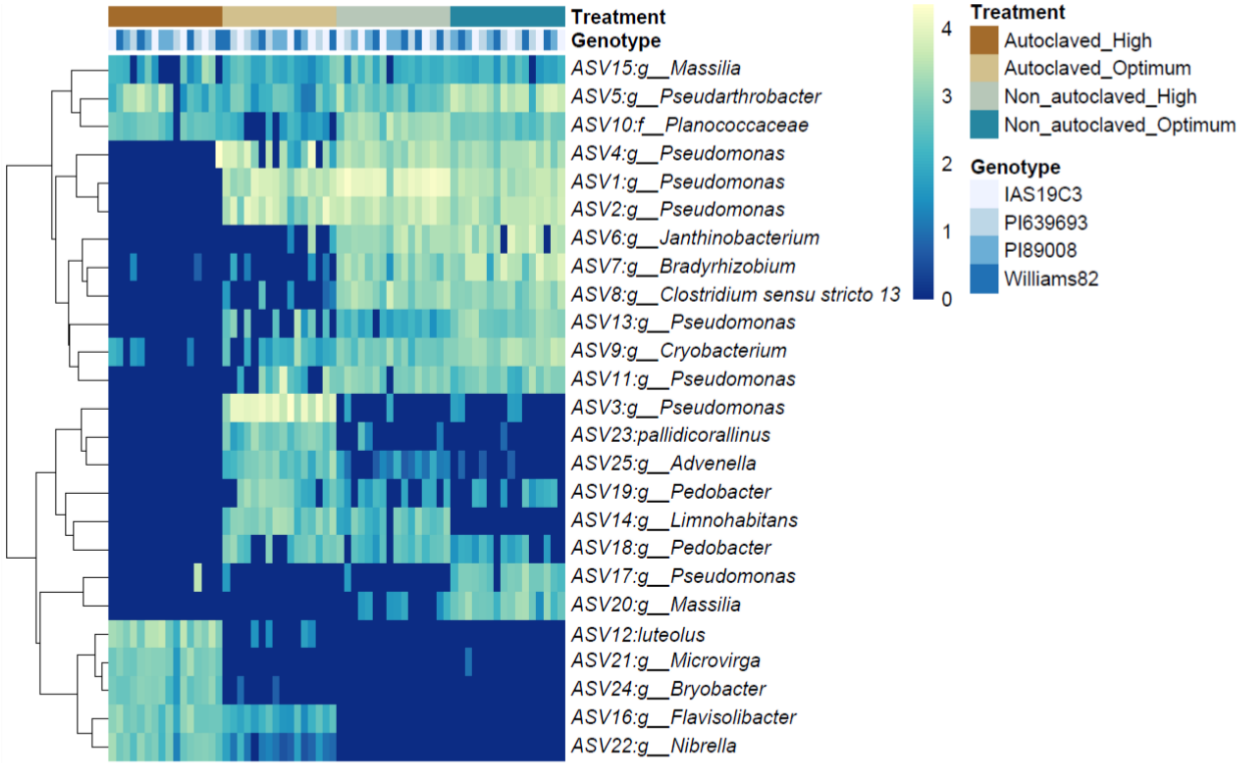

**Fig. S3**. **Rhizosphere-associated bacterial abundance was different between treatments**. Heat map of the top 25 ASVs showing different clusters derived from log transformation abundances of the bacterial lineages.

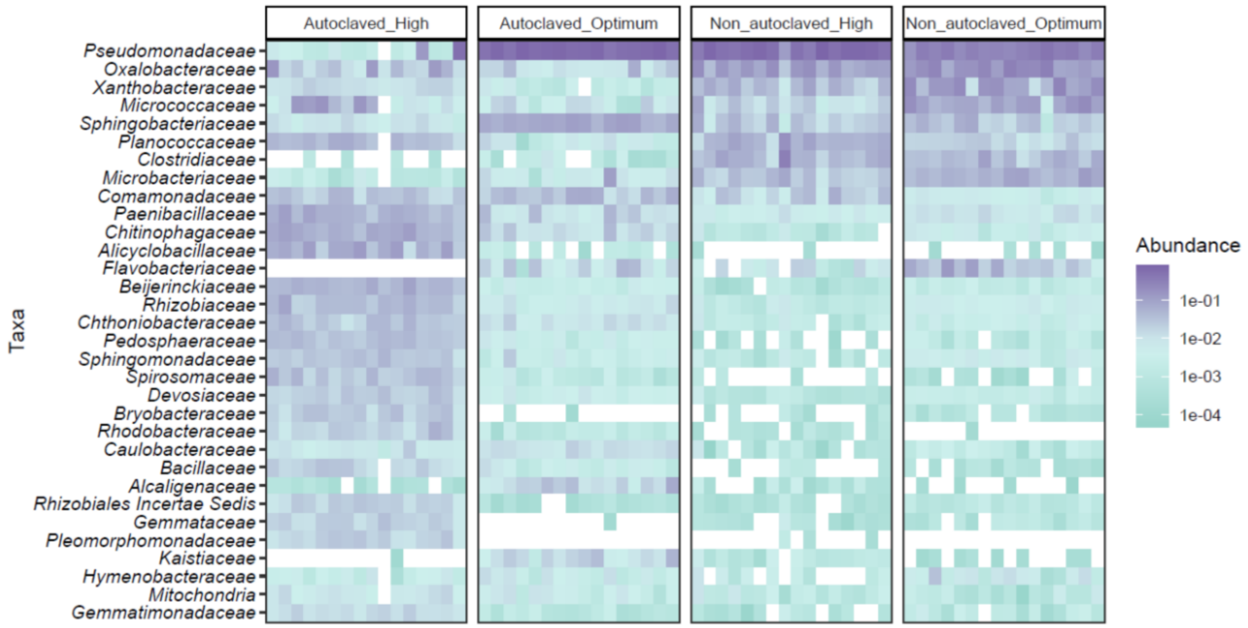

**Fig. S4**. **Heat map of differentially abundant bacterial taxa at family level across different treatments**. The heat map represents Log2fold expression. The violet and turquoise colors represent relatively high and low abundant levels, respectively.

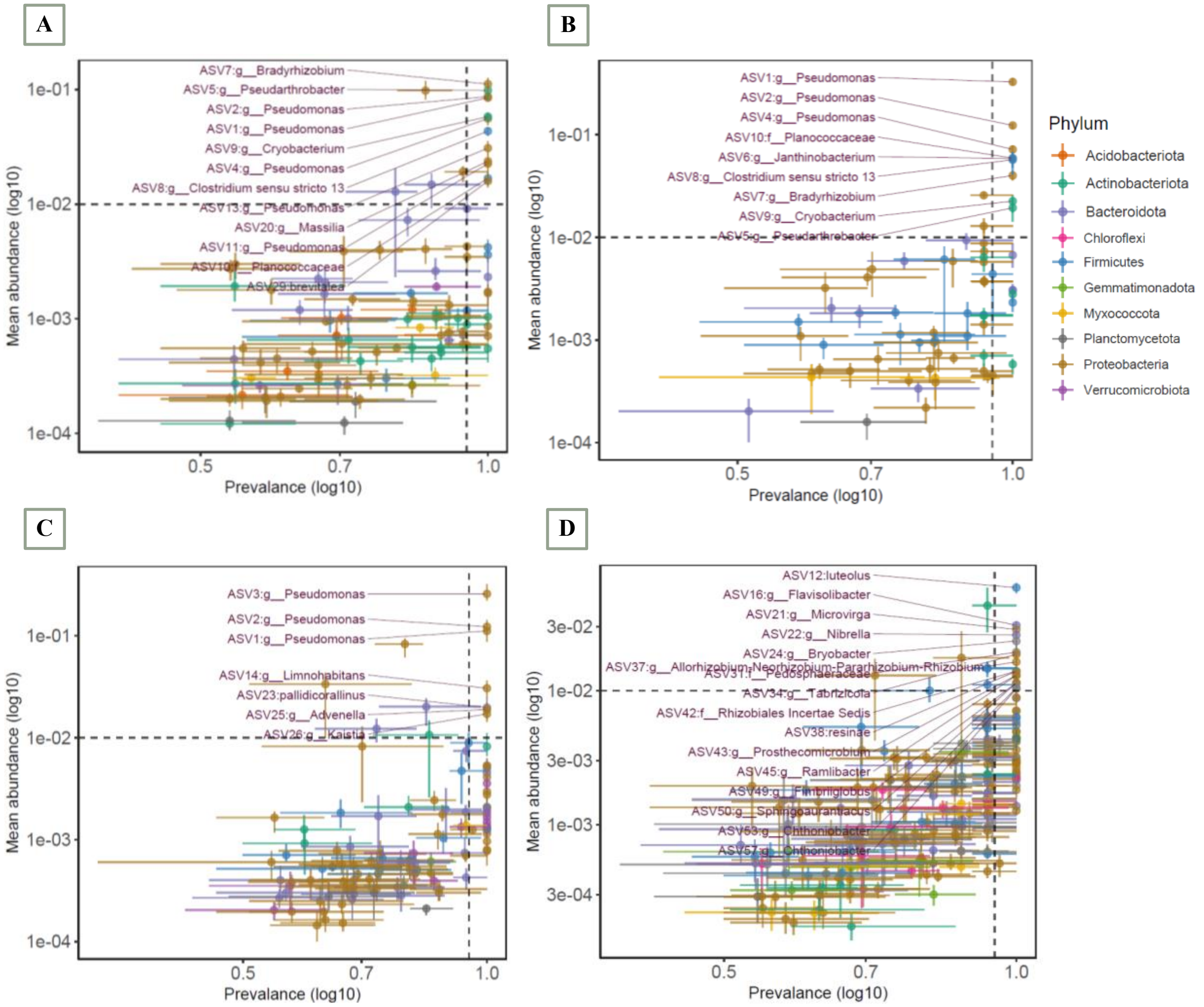

**Fig. S5**. Mean abundance-prevalence plots for bacterial taxa in **A.** non-autoclaved soil with optimum temperature (NO), **B.** non-autoclaved soil with high temperature (NH), **C.** autoclaved soil with optimum temperature (AO), and **D.** autoclaved soil with high temperature (AH). Mean abundance, mean prevalence, and upper and lower confidence interval for each ASV was calculated by random sub-sampling. Detection threshold is at abundance prevalence of 0.5 and detection level of 0.05.

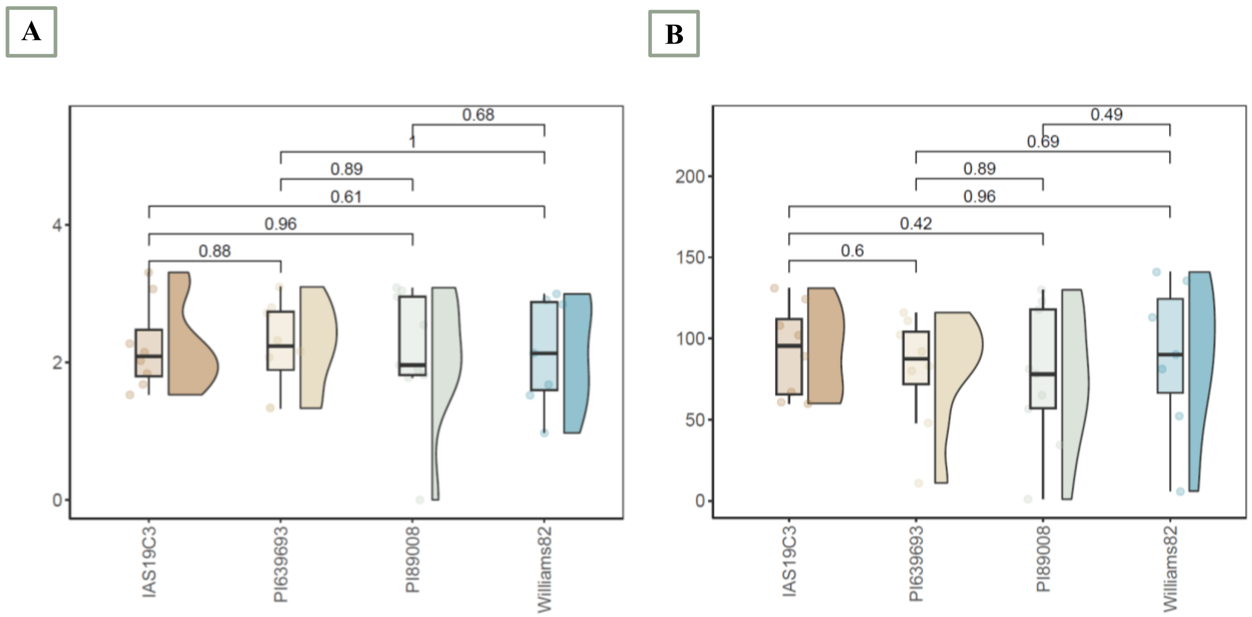

**Fig. S6**. **Alpha diversity measurements within the fungal communities associated with rhizosphere samples displayed no significant differences between the genotypes**. Estimated **A.** Shannon index and **B.** observed ASV richness in the fungal communities associated with the four soybean genotypes across four different treatments shown with ± SE and pairwise wilcox comparisons.

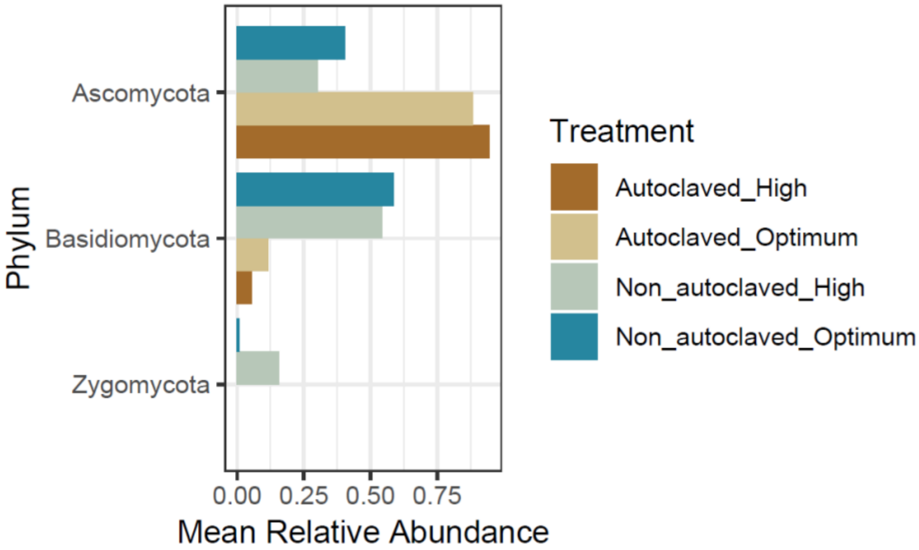

**Fig. S7** Differential taxonomic analysis presenting relative abundance of fungal taxa at the phylum level showed differences in distribution patterns between treatments.

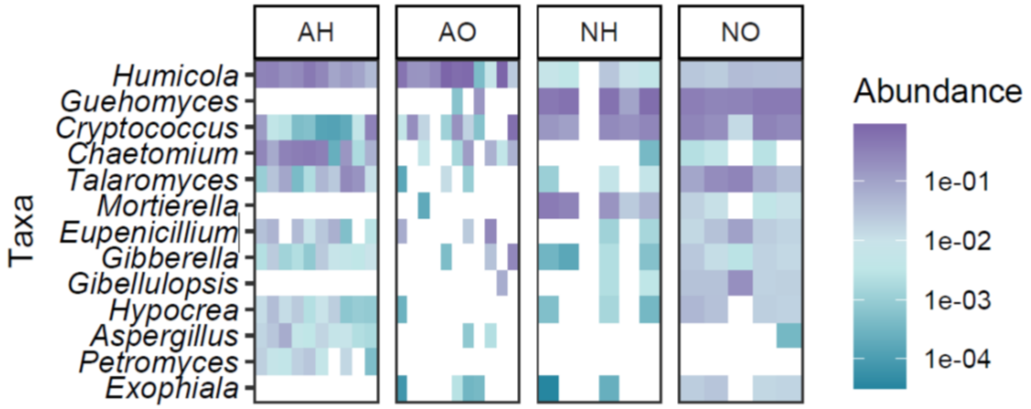

**Fig. S8**. **Heat map of differentially abundant fungal taxa at genus level across different treatments**. The heat map represents Log2fold expression. The violet color represents relatively high abundant, and turquoise color represents relatively low abundant level.

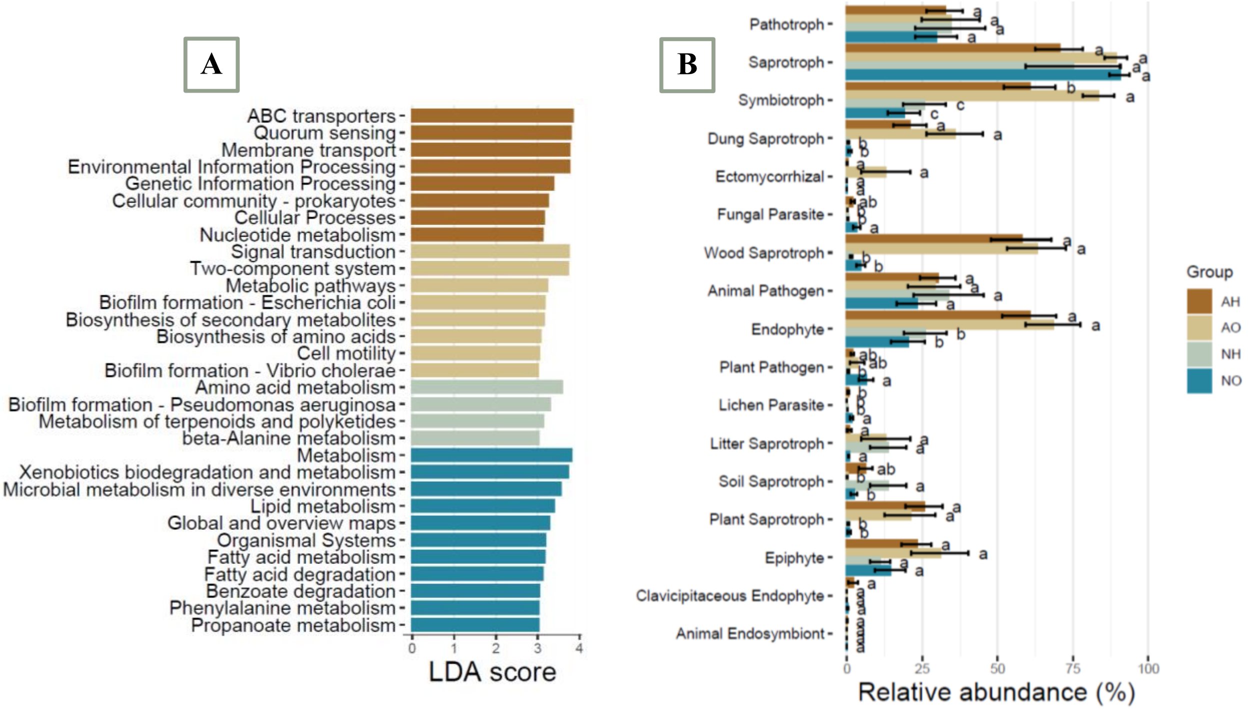

**Fig. S9**. Functional analysis of **A**. bacterial and **B**. fungal communities across different treatments.

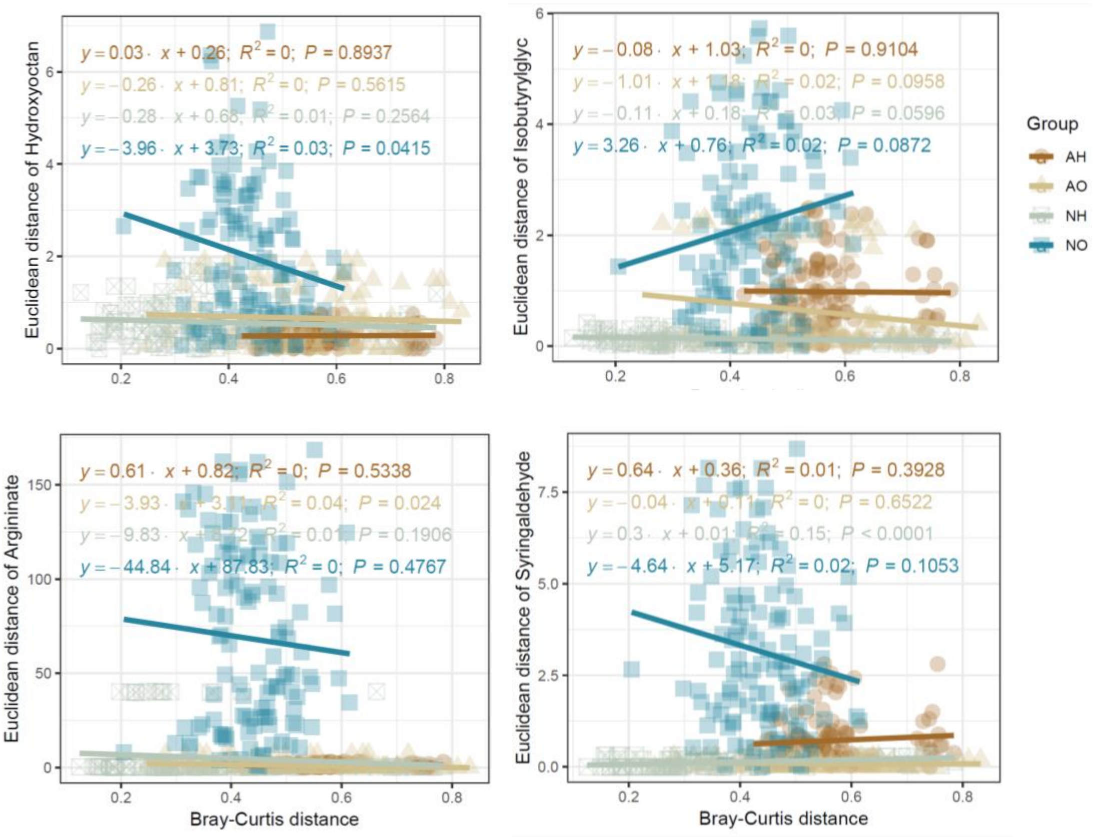

**Fig. S10.** Linear regression between root metabolites and rhizosphere bacterial communities. Points represent the mean value of the metabolites for soybean genotypes across different treatments.

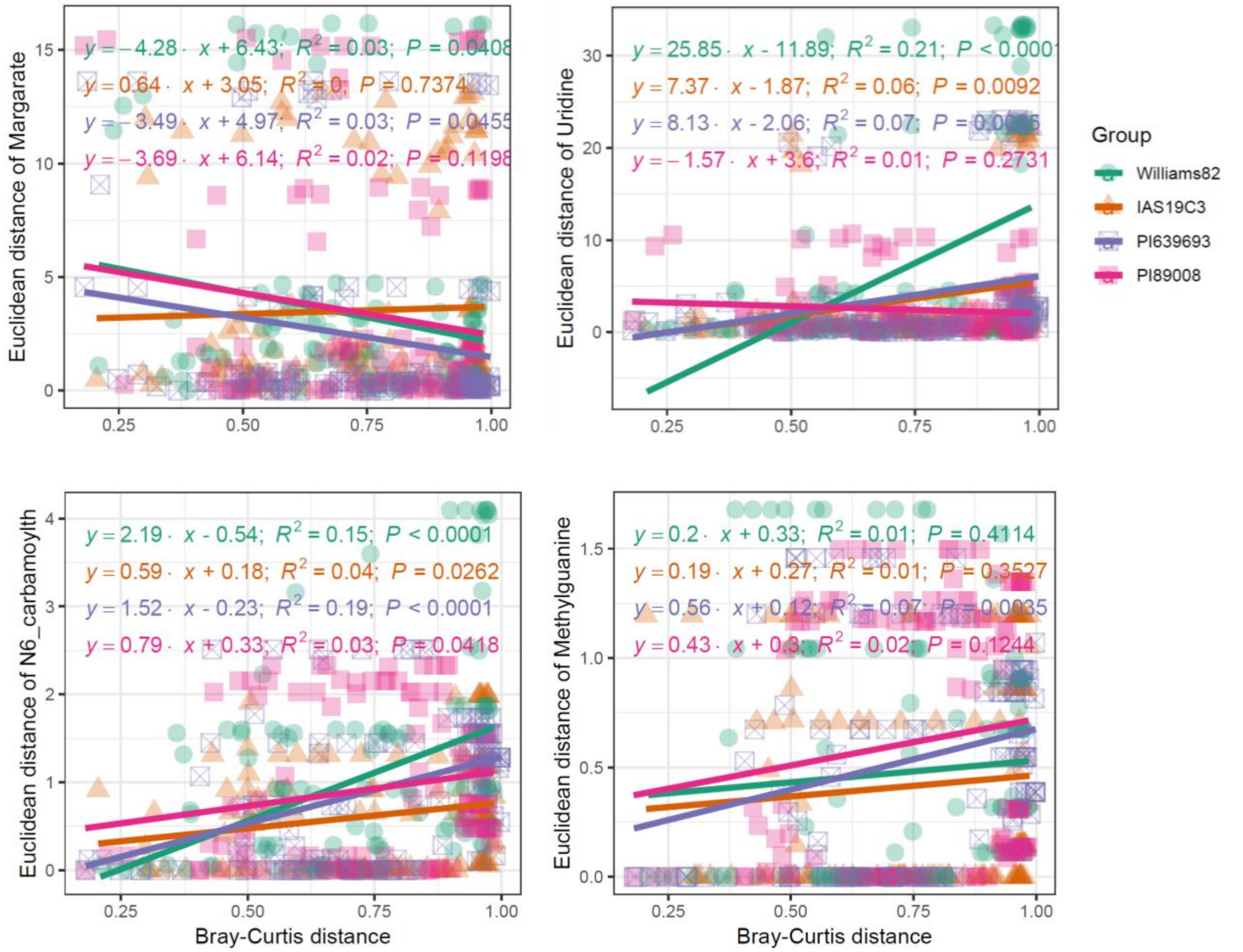

**Figure S11.** Linear regression between root metabolites and rhizosphere bacterial communities. Points represent the mean value of the genotype-responsive metabolites in soybean roots.

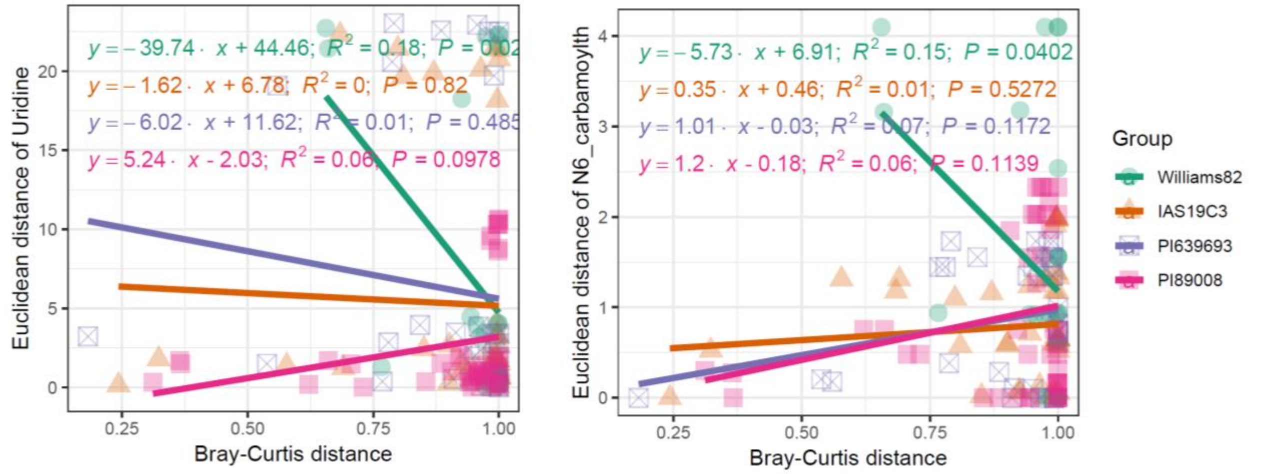

**Figure S12.** Linear regression between root metabolites and rhizosphere fungal communities. Points represent the mean value of the genotype-responsive metabolites in soybean roots.

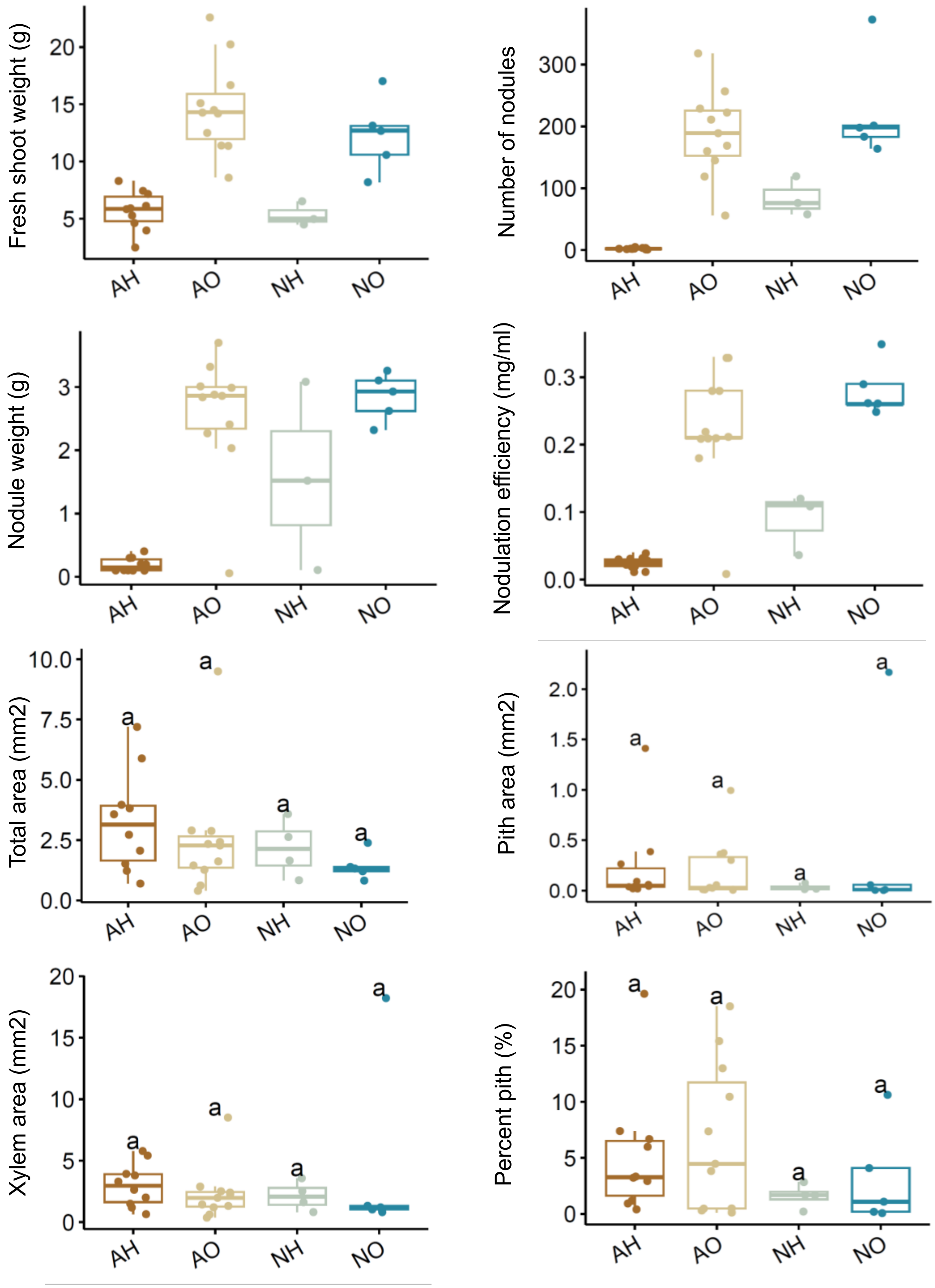

**Fig. S13**. The differential significance analysis of nodule and anatomical variables in the rhizosphere fungal communities of soybean submitted to different treatments.

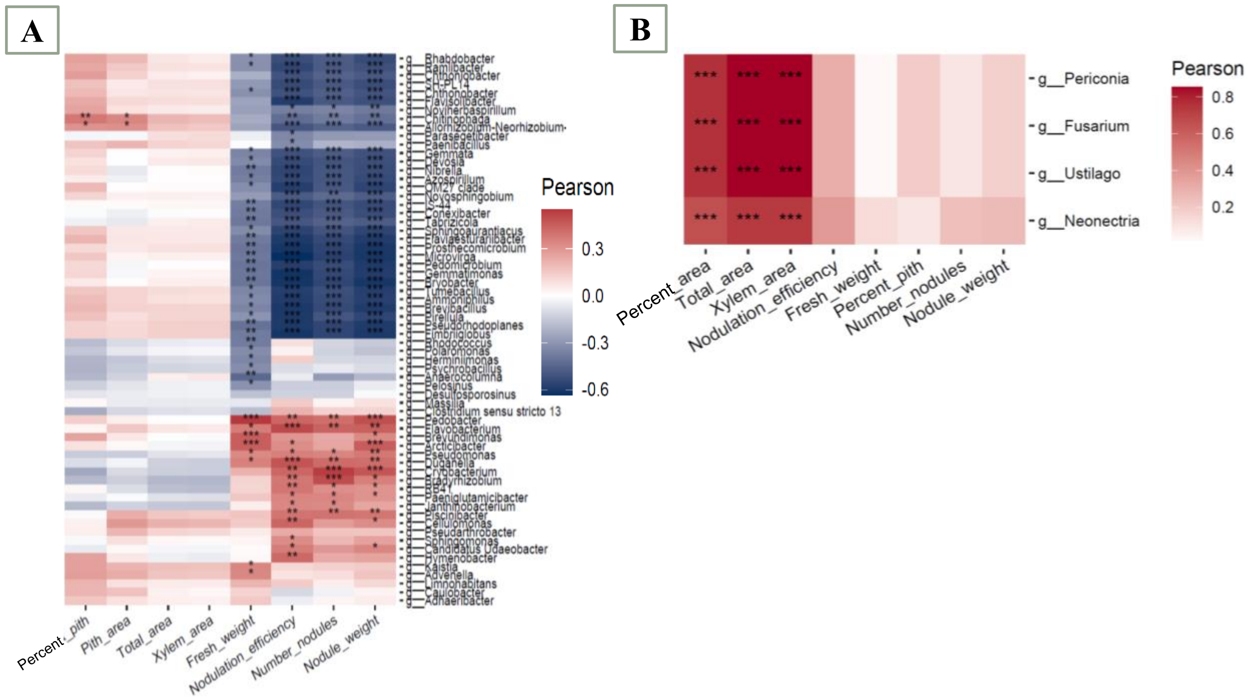

**Fig. S14**. Heatmap plots showing the correlations between the plant phenotypic variables and differential taxa in **A.** bacterial and **B.** fungal communities, respectively.
